# Social Dominance Reorganizes the Transcriptomic Neuropeptidome in a Highly Social Cichlid Fish

**DOI:** 10.1101/2024.11.07.622449

**Authors:** Isaac Miller-Crews, Hans A. Hofmann

## Abstract

Complex behavioral phenotypes, such as social status, emerge from the genome across biological levels, with many of the fundamental neural mechanisms shared across vertebrates. While various aspects of the brain have been implicated in modulating social behavior, cells of the preoptic area (POA) and hypothalamus are critical regulators of social dominance and applying cellular-resolution transcriptomic approaches allows for greater exploration of cellular dynamics. Yet, how complex gene networks function between and within cell types to regulate complex social behavior is still poorly understood. Importantly, when considering functionally relevant neuronal classes of genes such as neuropeptides, understanding the inherent complexity that emerges from the interaction of these genes in the transcriptomic neuropeptidome can provide unique insight into how social behavior is regulated. Here, we used single-nucleus RNA-sequencing in the hypothalamus and POA of a highly social cichlid fish, *Astatotilapia burtoni*, to understand the effect that social status has on distinct cellular-level transcriptomic profiles. Males of this species are well known for their highly plastic phenotypes related to social status, which allows for a hypothesis-driven approach. We demonstrate how social status manifests in changes of gene co-expression networks across neuronal populations, highlight known functional differences among AVP neuronal cell types, and implement a novel approach to relate how social dominance translates to a more highly integrated transcriptomic neuropeptidome. Taken together, this research provides insights into how gene expression networks that modulate social behavior, including neuropeptide networks, function at the cellular level.

## INTRODUCTION

Many of the neural mechanisms that regulate social behavior are shared across vertebrates, due either to evolutionary conservation or to convergence (for review, see (1, 2)). Numerous brain regions and neuron populations associated with different aspects of social behavior have been identified by their activity patterns (e.g., immediate-early gene, IEG, expression) and connectivity (e.g., tract tracing) (1, 3, 4). In recent years, open-ended genome-scale approaches, such as bulk or single-cell RNA-sequencing, have greatly expanded the scope of social neuroscience by uncovering the highly integrated and multifaceted nature of behavioral regulation by the brain (5–7). These advances also suggest an important and previously underappreciated role of glia cells in social behavior (8). However, how all these biological systems work together within and between specific cell types in relevant brain regions to encode and regulate complex social behavior is still poorly understood.

The hypothalamus and preoptic area (POA) are deeply conserved brain regions across vertebrates (9) and serve as key neural nodes in regulating basic homeostatic processes such as energy balance, reproduction, stress physiology and many other functions (10–12), including regulating social behavior (13, 14). The hypothalamus is a highly complex neural region, with multiple distinguishable subdivisions assigned by neuron morphology and function (15). In mammals, for example, the ventromedial hypothalamic nucleus (VMH) is well characterized for regulating aggression (16), while sex differences in the anteroventral periventricular nucleus (AVPV) can drive differences in parental behavior (17). Even though the homology relationships of these subdivisions across vertebrates are not always clear, studies in non-mammalian vertebrates suggest similar functions (18). Recent single-cell RNA-sequencing studies in the hypothalamus of mice have demonstrated the cellular complexity of the region (19–21) and how its cellular inventory is likely driving the functional differences in other brain regions (22). Due to the high degree of cellular heterogeneity of functionally distinct neurons compared to other brain regions (e.g., hippocampus, neocortex, cerebellum, etc.), structure-function relationships are often difficult to establish in the hypothalamus and POA (20, 23).

As a consequence of both the structural and peptidergic complexity of the hypothalamus/POA, a comprehensive and integrative understanding of the mechanisms through which this brain region regulates social behavior has been elusive. The situation is further complicated by the observation that hypothalamus and POA are a major source of neuropeptides. In fact, neuropeptides and the neurosecretory cells that produce them date back to the urmetazoan, the last common ancestor of animals, and possibly played a major role in the evolution of neurons (24, 25). Neuropeptides are enzymatically cleaved in the brain from larger precursor proteins (preprohormones) and often act as neuromodulators or neurohormones throughout the brain and, upon secretion from the brain, the entire organism (26, 27). Even small neuropeptides are known to coordinate a plethora of behavioral and physiological processes (28). And numerous neuropeptide signaling pathways have been implicated in specific homeostatic processes (e.g., energy balance: leptin, grehlin (29); reproduction: gonadotropin-releasing hormone (*GnRH 1*), gonadotropin-inhibiting hormone, kisspeptin (30); stress axis: corticotrophin-releasing factor (31); somatic growth: growth hormone (*GH*), somatostatin (*SST*) (32)). In mammals, over 130 peptide encoding genes have been identified, the vast majority of which are expressed in the hypothalamus and POA (33–35), and many also have important, evolutionarily conserved roles in regulating different aspects of social behavior (1). These genes usually encode at least two, and often more, preprohormones, affording peptidergic cells a staggering amount of combinatorial complexity and even more so when two or more of these genes are co-expressed (27, 28, 36). Therefore, to understand the role that neuropeptides play in regulating social behavior requires a systems-level perspective that examines the entirety of neuropeptide gene expression and co-expression within and across cells (37–40), what we call here the transcriptomic neuropeptidome.

The role of the hypothalamus and POA in regulating social dominance has been examined in detail in Burton’s Mouthbrooder cichlid, *Astatotilapia burtoni,* a highly social fish that has become an important model system in social neuroscience (41–43). Males of this species are characterized by highly-plastic dynamic phenotypes associated with social dominance status with well-established differences in neural gene expression patterns and neuron morphology (for review, see (44)). So, while all adult males are considered sexually mature, there is tight linkage between social status and reproductive state, with dominant males behaviorally suppressing reproduction in subordinate males, and with social status a transient phenotype dependent on the social structure. Importantly, *A. burtoni* easily allows for consistent engineering of individuals in distinct social status that serve to represent both a phenotype and social role, which have critical consequences on health and fitness. There has been considerable research on the relationship of social status and peptidergic signaling in the hypothalamus and POA in *A. burtoni* (45, 46). For example, both *GnRHI+* and *SST+* neurons are larger in size and have increased expression in dominant compared to subordinate males (32, 47). Likewise, three morphological subpopulations of vasopressin (*avp*) neurons, parvo-, magno-, and giganto-cellular vary by social status of both males (48) and females (49). Specifically, gigantocelluar *avp+* neurons have higher *avp+* expression in dominant males while parvocellular *avp+* neurons show the opposite effect. Even though these cells are morphologically distinct, and appear to have unique responses to social status, it remains unknown what differences exist at other biological levels, such as the transcriptome. Several other neuropeptide systems have also been implicated in the regulation of social dominance in *A. burtoni* (for review see (41, 43)). To compare across these neuropeptide systems, O’Connell & Hofmann (2012b) integrated behavior, physiology, and bulk transcriptomics of the POA in dominant and subordinate *A. burtoni* males demonstrating that several key pathways may serve as conduits for social plasticity across multiple levels (50). This suggests that environmentally induced changes at one level of biological organization do not simply affect changes of similar magnitude at other levels. However, previous studies lacked the cellular resolution that is necessary to match functionally important molecular pathways to specific cell types.

In the present study, we use single-nucleus RNA-Seq (snRNA-seq) to analyze the hypothalamus and POA of dominant and subordinate A. burtoni males. We investigated how differences in social status are linked to distinct transcriptomic profiles at the cellular level across three main hypotheses. Even though prior research has shown that the population size of at least some peptidergic cell types can vary by social status, most research has focused on quantitative differences in the expression levels of peptide-encoding genes. We first hypothesized that, overall, dominant and subordinate males would possess similar inventories of neuronal and non-neuronal cell types to maintain basic hypothalamic function. Second, we hypothesized that morphologically distinct and socially responsive subpopulations of *avp+* neurons (as demonstrated by Greenwood et al., 2008) represent unique transcriptomic nuclei clusters. Third, given the role neuropeptides play in regulating social status in *A. burtoni* and across vertebrates, we hypothesized that preprohormone expression and co-expression patterns across the transcriptomic neuropeptidome would reflect social status in specific cell types; specifically, that dominant males would display a more tightly organized transcriptomic neuropeptidome to support their compared to subordinate males much more complex behavioral repertoire. Together, our analyses aim to provide a comprehensive accounting of the transcriptomic mechanisms through will social dominance status emerges from the hypothalamus in a cell type-specific manner.

## RESULTS

### Cell type identification of sequenced hypothalamus nuclei

We identified 26,777 single-cell transcriptomes across all samples and successfully demultiplexed the three expected genotypes per sample pool between dominant and subordinate *A. burtoni* males, providing biological replicates for downstream analyses (Fig 1a, 1b). Cell-type calling demonstrated consistency of cell types between dominant and subordinate males and biological replicates (Fig 1c; Fig S1a) for six major hypothalamic neural cell categories: excitatory neurons, inhibitory neurons, oligodendrocytes, astrocytes, vascular cells, and cells of the *pars tuberalis* (21).

**Figure 1:**
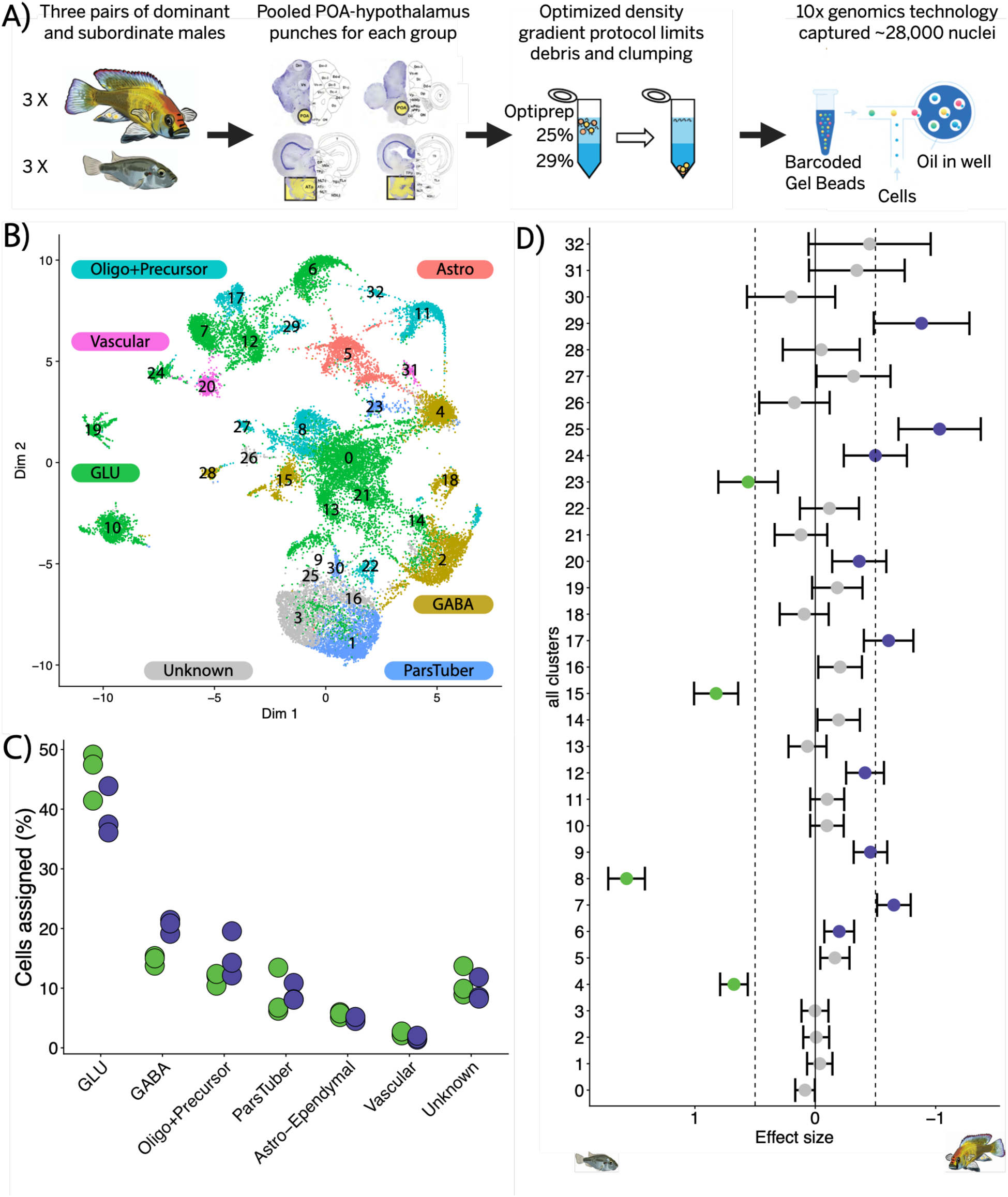
Experimental overview for snRNA-seq pipeline and cell type identification. A) Three pairs of dominant (top) and subordinate male (bottom) *A. burtoni* were collected and the POA-hypothalamus (highlighted in yellow) were pooled for each group prior to single nuclei isolation and snRNA-seq with 10x genomics. Fish illustrations courtesy of Dr. Caitlin Friesen. Brain images from (116). Barcode image courtesy of 10x Genomics. B) UMAP projection of single nuclei data with statistical clusters numbered and cell type assignments color-coded with labels. C) Cell type identification resulted in roughly equal percentage cells assigned across all genotypes for dominant (purple) and subordinate (green) males. D) Forest plot depicting effect of social status on percentage of nuclei in each cluster and 95% confidence interval with color denoting significant difference (p-value < 0.01) between dominant (purple) and subordinate (green). Dashed lines indicate absolute effect size of 0.5.

One way in which social status differences may manifest at the cellular level is through differences in the number of certain cell types. We therefore wanted to test whether there was any difference in the proportions of cell subtypes due to social status. We compared the proportion of nuclei clusters by social states with a binomial GLM (for detailed statistics see: Dataset S1). Of all nuclei clusters, thirteen exhibited biased proportions of nuclei towards either dominant (clusters 6, 7, 9, 12, 17, 20, 24, 25, 29) or subordinate males (clusters 4, 8, 15, 23) (Fig 1d; Fig S1b). Of the four clusters with a dominant bias, three GABAergic (cluster 4 and 15), one was annotated as pars tuberalis (cluster 23), and one as oligodendrocytes (cluster 8). Of the nine clusters with a dominant bias, five were glutamatergic neurons (clusters 6, 7, 9, 12, and 24), one contained vascular cells (cluster 20), and two oligodendrocytes (cluster 17 and 29), with one cluster lacking any annotation (“unknown:” cluster 25).

Since only a few non-neuronal cell types showed proportional differences between social states, we next asked whether gene expression patterns differed between dominant and subordinate males in those cell types. For each of the broad nuclei clusters, we identified numerous differentially expressed genes (DEGs) between dominant and subordinate males at the cell-type specific sub-cluster level (Fig S2a; Dataset S2). We found that the vast majority of status-linked DEGs between clusters were due to changes in the intensity of gene expression per cell, rather than changes in the proportion of cells expressing that gene. Of those, five genes emerged as dominant-biased fraction-driven DEGs in almost every nuclei cluster, including *cga*, *gh1, prl, oxt,* and *gnrh1*, potentially representing a statistical artifact (Fig S2b). To mitigate this issue, we used in all subsequent analyses dynamic thresholds corresponding to how broadly expressed a gene was when testing for differences in the proportion of nuclei that express those genes. Yet the presence of large numbers of DEGs across all nuclei clusters, not just neuronal clusters, demonstrates that all cell types reflect social status to some extent.

### Neuron transcriptomic networks and social dominance

Because a variety of hypothalamic neuron types have long been implicated in regulating social status across vertebrates, we next examined neuronal cell populations in some detail. First, graph-based clustering of neurons revealed highly structured subpopulations of excitatory (i.e., glutamatergic) and inhibitory (i.e., GABAergic) neurons (Fig 2a; Fig S1c). We then compared the proportion of neuron clusters between dominant and subordinate males with a binomial GLM that included individual genotypes (for detailed statistics see: Dataset S1). Thirteen of these neuron clusters were found to have biased proportion of nuclei (FDR p-value < 0.01) favoring either dominant (neuron clusters 2, 5, 6, 8, 14, 15, 16, and 19) or subordinate males (neuron clusters 1, 7, 9, 11, and 18) (Fig 2b; Fig S1d).

**Figure 2:**
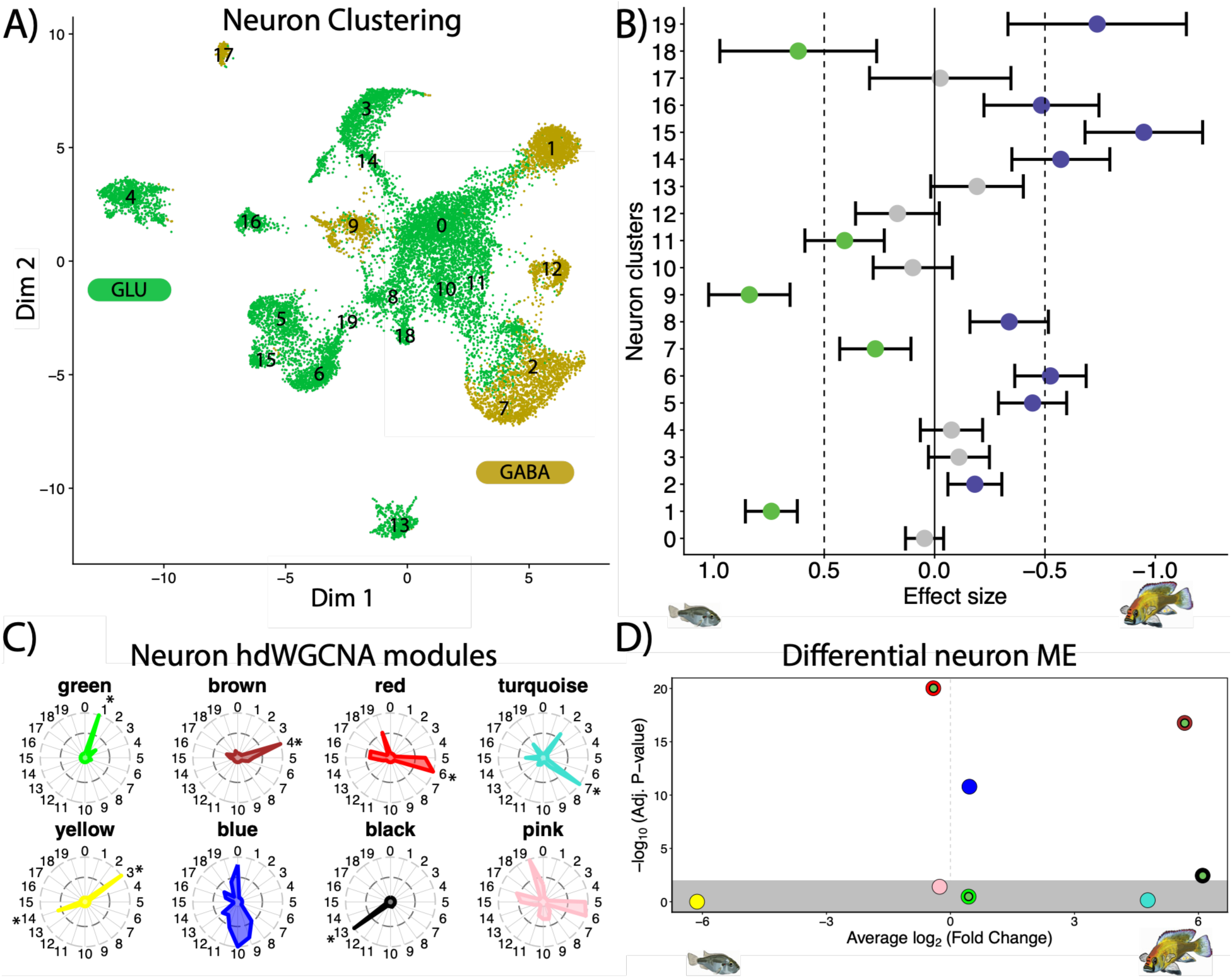
Neuron gene network analysis. A) UMAP projection for all neuron nuclei with statistical clusters numbered and color-coded with labels into excitatory (GLU = green) and inhibitory (GABA = yellow) neurons. B) Forest plot depicting effect of social status on percentage of nuclei in each neuron cluster and 95% confidence interval with color denoting significant difference (p-value < 0.01) between dominant (purple) and subordinate (green). Dashed lines indicate effect size of 0.5. C) Implementation of single-cell optimized weighted gene co-expression network analysis, hdWGCNA, identified eight gene expression modules. hdWGCNA module eigengene (ME) values across neuronal clusters illustrate how most gene co-expression networks are expressed within certain neuron clusters. * indicate specific module-neuron cluster pairs (z-score > 2). D) Gene modules show significant differential module eigengene expression by social status among all cells. For modules with specific module-neuron cluster pairs, inner circle colors indicates presence of significant (FDR < 0.05) difference in average MEG expression by social status within that specific neuron cluster (green = subordinate, purple = dominant).

Next, we assessed the degree of differential expression within neuron clusters by social status. Similar to the broad nuclei clusters, each neuronal clusters displayed DEGs biased towards either dominant and subordinate males (Fig S3a), with fewer DEGs in smaller clusters due to decreased statistical power with fewer nuclei available for testing (Dataset S2). Likewise, we identified the same core group of 5 dominant-biased genes in all clusters as was the case in the broad cluster DEG analysis, with the exception of neuron cluster 11 being the only cluster to lack a difference in *oxt* expression by social status. As an example, beyond the core genes differentially expressed in every cell type minus *oxt*, neuron cluster 11 has three subordinate-biased genes (*grid1a, utrn, tmem254*) and five dominant-biased genes (*eef2b*, *pomc, lhb, si:dkey-96f10.1, pfkfb1*).

Due to the known role of specific neuronal cell types in regulating social behavior, we hypothesized that there would be differential activity patterns between neuron clusters reflecting differences in cell subtype. While this experiment was not designed to directly test differences in activity level, we did note behavioral differences related to social status prior to sample collection that would likely lead to changes in neuron activity, along with systemic differences due to differences in behavioral state between dominant and subordinate males (Supplemental §1). We compared differences in the proportion of immediate early genes (IEG) expressing neurons across a panel of known IEGs (*egr1, npas4, fosb, fosab*) across neuron clusters and between social states (Fig 3b). Neuron cluster 11 demonstrated the highest proportion of IEG expressing nuclei at 48.8% ± 12.3. Only neuron cluster 10 was identified to have significantly higher proportion of IEG expressing nuclei in subordinate males (*t*(2.19) = −4.07, FDR = 0.054), with no neuron cluster displaying a dominant-bias.

**Figure 3:**
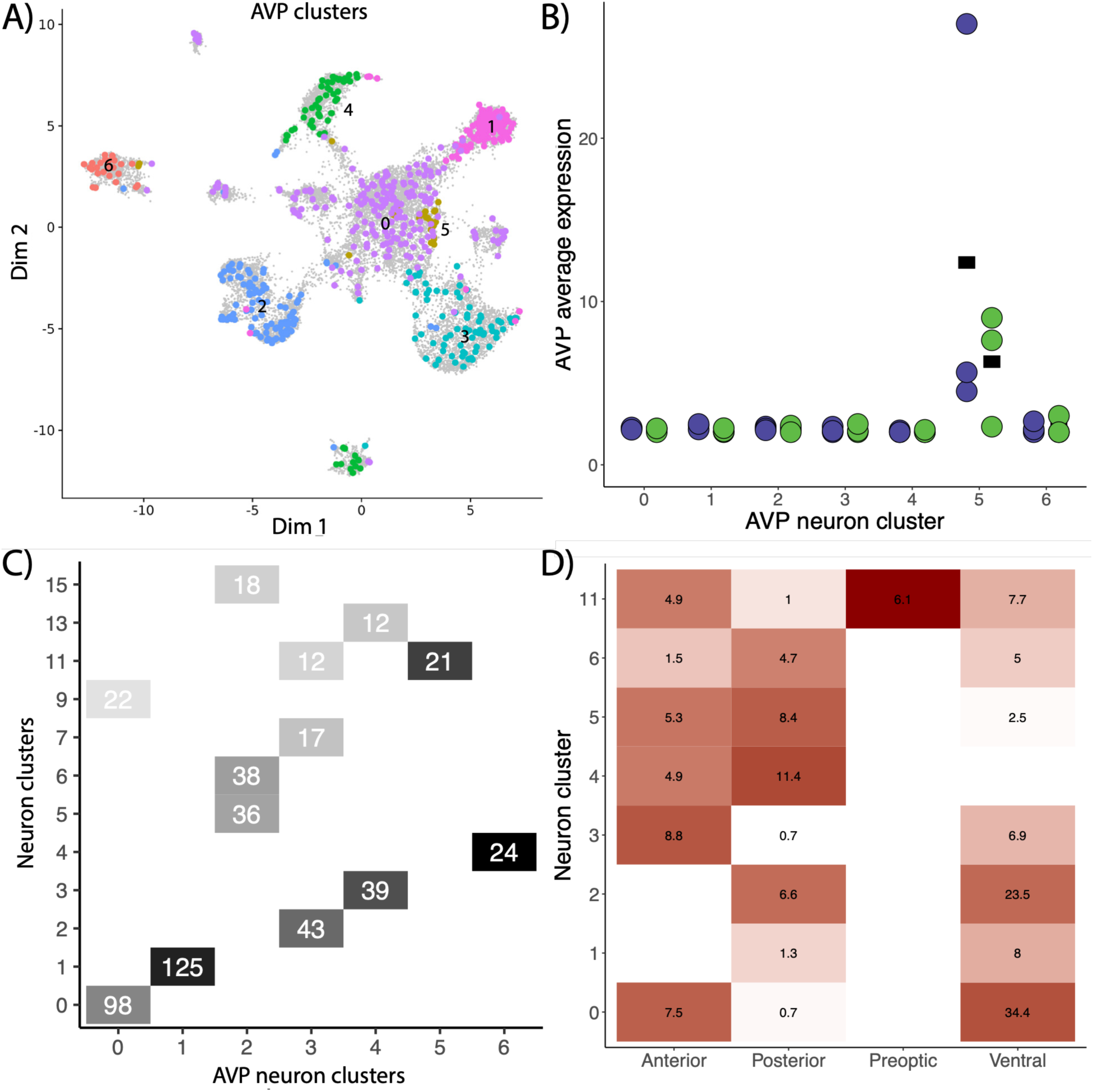
AVP expressing neuron subpopulations. A) Projection on neuron UMAP of all AVP expressing neurons colored and labelled by the five statistical subpopulation clusters. B) Average AVP expression of each AVP neuron statistical clusters for each of the dominant (purple) and subordinate (green) male genotypes, with black bars indicating mean. C) Comparison of overlap between nuclei assigned to AVP neuron clusters (x-axis) and neuron clusters (y-axis). Numbers indicate the number of nuclei overlapping with colors representing the proportion of AVP neuron cluster assigned to each neuron cluster with a threshold requirement of at least 10% and 10 nuclei overlapping. D) Spatial prediction across broad hypothalamic zones for the subset of neuron clusters (y-axis) with the highest proportion of assigned nuclei from each AVP neuron cluster. Heatmap displays the proportion of weighted permutation z-scores within a zone for mapping to HypoMap regions, collapsed into broad hypothalamic spatial zones with known AVPcells. Values indicate higher confidence in spatial assignments after permutation filtering and region weighting.

To gain a better understanding of which parts of the POA and hypothalamus these neuronal nuclei represent, we used the HypoMap database (21) to assign each nucleus to a putative preoptic or hypothalamic sub-region (Supplemental §3). Briefly, we first assigned cichlid orthologs to the marker genes of the HypoMap clusters (Fig S4a). Next, we assigned Hypomap clusters to spatial locations, and subsequent regional zones, using the proportion of nuclei from each cluster with known spatial origins (Fig S4b). We then calculated a weighted permutation z-score of Spearman correlations between marker genes for each pair of HypoMap clusters and neuronal clusters from the present study (Fig S5a). This allowed us to estimate spatial origin within regional zones for each neuronal cluster (Fig S5b). This approach revealed that the majority of neuronal nuclei in our dataset corresponded to specific preoptic and hypothalamic subregions. Of note, neuron cluster 11was the highest scoring cluster for the preoptic zone and therefore our best candidate for the POA.

We next asked whether and how gene co-expression networks might vary by social status and across neuronal sub-types. To do this, we employed hdWGCNA by generating metacell profiles for each of the six demultiplexed genotypes (see above). We found seven gene co-expression modules that were expressed across neurons, although not evenly across neuron clusters (Fig 2c; Fig S6a). Differential module eigengene (MEG) expression analysis revealed several gene modules that were significantly differentially expressed (FDR p < 0.01) by social status (Fig 2d), with three modules significantly overexpressed in dominant males (brown, blue, and black modules) and one module significantly overexpressed in subordinate males (red module), suggesting that across all neuronal nuclei the activity of certain gene networks is social status-dependent.

To better understand the relationship between modules and cell-type, we then asked whether social status differences in MEG expression were cell type-or cell cluster-specific. We first identified the module that was most representative for a given cluster (i.e., its MEG expression was highest in that cluster) to generate enriched module-cluster pairs (Fig 2c). For each module, we discovered a neuron cluster (or clusters) with the most specific expression for a given module: 1-green; 3,14-yellow; 4-brown; 6-red; 7-turquoise; 13-black (neuron cluster-maximum module). We then assessed MEG expression differences between dominant and subordinate males within each specific module-neuron cluster pair using linear models on pseudo-bulk MEG means of each individual genotype within each cluster, weighted by the relative number of nuclei for that genotype in that cluster. Of all the module-neuron cluster pairs listed above, four showed significantly higher MEG expression (FDR p < 0.05) in subordinates: 1-green; 4-brown; 6-red; 13-black (Fig 2d; Fig S6b). Besides the red module, these results were in contrast to the neuron-wide differential MEG expression analysis, indicating that compositional shifts (e.g., biases in the number of nuclei per cluster) may play an important role across all neurons.

To identify putative functions of these modules we used GO enrichment analysis (Dataset S3; Fig S6c), which revealed significantly enriched GO terms related to: *system development* in the brown module; *cell-cell adhesion* in the blue module; and *cell-cell adhesion* and *cell communication* in the pink module. For modules with many enriched GO terms, identified terms summarized as: *regulation of development and cell differentiation*, *protein dephosphorylation*, and *actin filament organization* in the black module; *gene expression*, *peptide processing*, and *mitochondrial ATP synthesis* in the green module; and *cell communication*, *synaptic signaling*, and *ion transport* in the turquoise modules (Fig S6d). We found no significantly enriched GO terms in the red or yellow modules (Dataset S3).

### Neurons expressing arginine vasopressin (AVP) pre-prohormone mRNA

Previous research in *A. burtoni* identified three preoptic AVP-positive neuron populations that varied in size (representing the parvo-, magno-, and gigantocellular neurons) and were regulated differently by social status (48). We reasoned that these three morphologically characterized *avp+* neuron populations may also emerge from our single cell data. To test this hypothesis, we first identified all neurons that expressed *avp+* preprohormone mRNA, resulting in 363 and 270 nuclei for dominant and subordinate males, respectively. Graphical clustering of these *avp+* neurons discerned seven clusters that were present in all six genotypes (Fig. 3a). Next, we asked whether *avp* expression levels differed across *avp+* neuron clusters and social status using a gamma glm (Fig. 3b). The Type III analysis of deviance found a main effect of cluster (χ² = 102.71, df = 6, p < 2×10⁻¹⁶), with no significant effects of status (p = 0.892) or status-by-cluster interaction (p = 0.504). Using Tukey *post hoc* comparisons, we identified *avp+* neuron cluster 5 as having significantly higher *avp* expression compared to all other clusters (all p<0.0001), with no other significant differences amongst clusters (p=1). Importantly, *avp* expression did not differ by social status in any of these clusters (all FDR p > 0.13).

Given the high *avp* expression in *avp* neuron cluster 5, it is noteworthy that the broad neuron cluster it corresponded to (neuron cluster 11) also expressed high levels of other neuropeptide genes such as *oxt* (Fig S7b). Additionally, since this same neuron cluster 11 contained the highest proportion of IEG expressing nuclei among all neuron clusters, we assessed social status-dependent activation differences. Specifically, we compared average IEG expression between dominant and subordinate genotypes across a panel of known IEGs (*egr1, npas4, fosb, fosab*) for each of the *avp+* neuron clusters and discovered that *egr1* expression in *avp+* neuron cluster 1 was higher in dominant males, while expression of *npas4, fosb, and fosab* in *avp+* neuron cluster 5 was higher in subordinate males (Fig S7c; Fig S7d). Although these results should be considered preliminary, they do suggest that *avp+* neuron clusters 1 and 5 may play specific roles in dominant and subordinate males, respectively.

As subpopulations of *avp+* neurons are known to be spatially dispersed across the teleost POA and hypothalamus (48, 51), we examined whether any of the *avp+* neuron clusters described above could be assigned to specific POA or hypothalamic subregions. Using a spatial location prediction score (See Supplemental §3) for the broader neuron clusters that best encompassed each *avp*+ neuron cluster, we compared the overlap of nuclei in each *avp+* neuron clusters to broader neuron clusters (Fig. 3c) and assigned the putative spatial location to *avp+* neuron clusters based on the broader neuron cluster that the majority of each *avp+* neuron cluster was part of (Fig. 3d). We found the strongest evidence for *avp+* neuron clusters 0, 1, 2, and 3 in the ventral hypothalamus, for *avp+* neuron cluster 4 in the anterior hypothalamus, for *avp+* neuron cluster 5 in the POA, and for *avp+* neuron cluster 6 in the posterior hypothalamus.

### Uncovering the transcriptomic neuropeptidome

To better understand the highly complex and combinatorial system of hypothalamic neuropeptides, many of which are known to regulate social behavior, we examined the entirety of neuropeptide gene expression and co-expression within and across cells, what we propose to call the transcriptomic neuropeptidome. For each neuron we counted the number of different neuropeptide genes expressed within each sample. For the 67 neuropeptide preprohormones we were able to annotate in our dataset, we found that the vast majority of neurons express at least one of them. Next, we compared the distribution of neuropeptides per neuron between dominant and subordinate males (Fig 4a). We found that the distribution of the number of neuropeptides per neuron was from the same distribution for: dominant males (AD = 0.874, p = 0.965), subordinate males (AD = 0.1.25, p = 0.788), and across all samples (AD = 2.31, p = 0.995). Similarly, we found no difference in the number of neuropeptides per neuron within dominant males (Poisson, β = −0.021 & 0.017, SE = 0.019 & 0.017, p = 0.259& 0.321) or within subordinate males (Poisson, β = −0.000 & −0.014, SE = 0.024 & 0.025, p = 0.988 & 0.590) indicating that the counts were consistent across individual genotypes of the same social status. However, we found that dominant male neurons have a significant 33% increase in neuropeptide genes expressed compared to subordinate male neurons (Poisson, β = −0.405, SE = 0.012, p < 2e-16).

**Figure 4:**
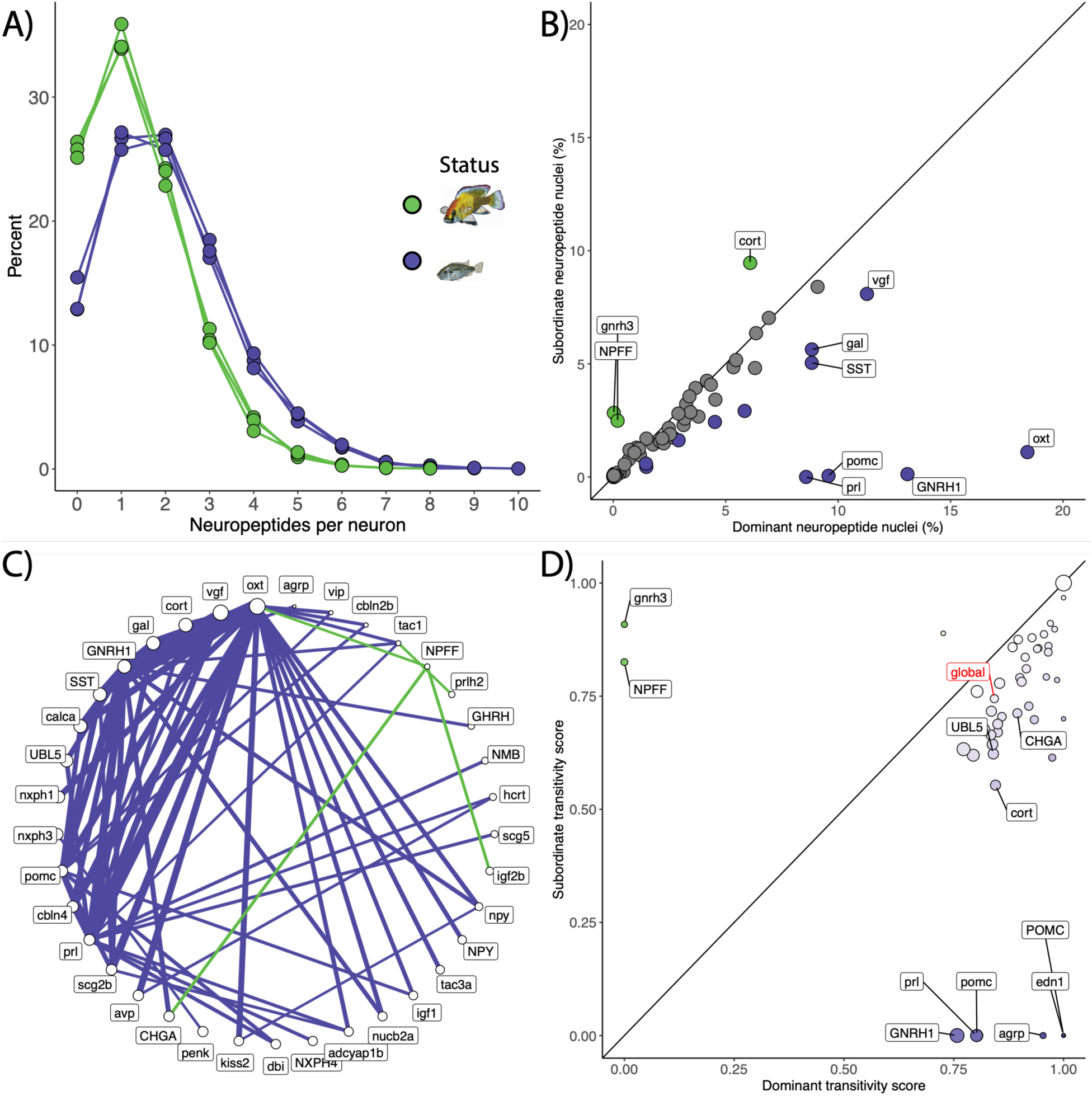
Uncovering the Transcriptomic Neuropeptidome. A) Neurons broadly express neuropeptide genes, dominant male (purple) neurons have a higher percentage of neuropeptide co-expression than subordinate (green). B) The average percent of neurons expressing each neuropeptide by social status with proportionally significantly different (Binomial GLM FDR p-value < 0.001) neuropeptides colored and labelled for dominate bias (purple) and subordinate bias (green). C) Significantly (FDR p-value < 0.05) different edges between the dominant (purple) and subordinate (green) neuropeptide gene co-expression network. D) Comparison of network clustering through transitivity by social status both global (red label) and locally (black label) for each neuropeptide gene, with higher value indicating greater shared connections. Dot size indicates the weighted node strength with higher values indicating more highly connected genes, and color indicates difference in node strength between dominant (purple) and subordinate (green) males.

To uncover which neuropeptide genes were differentially expressed in neurons of dominant and subordinate males, we compared the number of neurons expressing each neuropeptide to find neuropeptide genes that varied in expression by social status (Fig 4b). We found that the proportion of neuronal nuclei expressing specific neuropeptide genes was significantly different (FDR p <0.001) between dominant and subordinate males (for detailed stats on binomial GLM results see: Dataset S1): there were three neuropeptide encoding genes with a subordinate bias (*cort, NPFF, gnrh3*) and 12 with a dominant bias (*GNRH1, SST, agrp, gal, kiss2, npy, oxt, pomc, scg2b, vgf, vip, prl*).

Since neuropeptide gene co-expression was so widespread, we next quantified the difference across the transcriptomic neuropeptidome between social states. Using neuropeptide co-expression networks normalized by total nuclei counts, pairwise network similarity was higher among dominant than subordinate males (Dominant-Dominant: r = 0.919 ± 0.016 [mean ± sd]; Subordinate-Subordinate: r = 0.756 ± 0.015), and significantly more similar within social status than between (Dominant-Subordinate: r = 0.243 ± 0.019; Wilcoxon test P = 0.0004). This indicates that pairwise network similarity is three times greater within social status than between. PERMANOVA identified strong separation in co-presence networks related to social status (R^2^ = 0.857, p = 0.1), with significant differences in network variability between social states (beta dispersion: p = 0.001), suggesting that social status differences are reflected in both network structure and variability.

Because co-expression of neuropeptide genes was so common across preoptic and hypothalamic neurons, with only a few neuropeptide expressing cell populations displaying social status-dependent nuclei counts, we determined which pairs of neuropeptide genes, if any, differed in their co-expression by social status. We used neuropeptide gene co-expression networks to find the difference between individual dominant and subordinate genotypes for each pair of neuropeptide genes (Fig 4c; Fig S8) and found widespread significant increases (FDR p < 0.15) in neuropeptide co-expression in dominant males, with the exception of the significantly higher co-expression in subordinate males between *NPFF* and four other genes (*oxt, prlh2, igf2b, CHGA*). To assess whether neuropeptide co-expression networks differed in how closely clustered they were between social states we calculated a global transitivity score, a network metric that measures the probability that the neighbors of one node are also connected to each other (Fig 4d). Global transitivity of the neuropeptide gene co-expression networks indicated a higher, though not significant, average probability of shared connections among nodes in dominant compared to subordinate males (t(2.23)=3.38, p-value = 0.067). Likewise, we calculated local transitivity of each neuropeptide gene within the network to quantify how connected the neighbors of that gene were to each other. Of the 39 neuropeptide genes tested, three (*UBL5, CHGA, cort*) differed significantly by social status (FDR p < 0.05) and eight only had transitivity in either subordinate (*gnrh3, NPFF*) or dominant (*GNRH1, prl, pomc, agrp, POMC, edn1*) males. These results indicate that the expression of neuropeptide genes in the transcriptomic neuropeptidome was more tightly clustered in dominant males compared to subordinate males, suggesting increased gene co-expression network connectivity in the former. Of note, three neuropeptide genes with exclusive dominant-biased local transitivity also had high node strength (*gnrh1, prl, pomc*). In other words, dominant males not only contained neurons uniquely expressing these neuropeptide genes but these genes also were part of status-specific neuropeptide co-expression networks.

## DISCUSSION

In the present study, we examined how social status is reflected in the variation of cell type-specific transcriptomic profiles across the hypothalamus and POA. We successfully implemented single-nucleus RNA-seq in a non-traditional model organism to identify the transcriptomic changes associated with social dominance in the hypothalamus and POA of males in *A. burtoni*. In contrast to our first hypothesis, we observed distinct changes in the expression of neuron gene co-expression networks indicating that the brains of both dominant and subordinate males engage in active maintenance of discrete pathways. Following our second hypothesis, our approach was able to target specific known functional neural populations of *avp+* neurons to assess the impact of the social environment on gene expression. Finally, in relation to our third hypothesis, we demonstrated how the transcriptomic neuropeptidome changes with social state within cells, going beyond what these neuropeptide genes do independently, and that dominant males exhibit a more highly integrated transcriptomic neuropeptidome. Taken together, our results demonstrate the remarkable complexity and plasticity of cell type-specific transcriptomes in relation to social status.

### Neural transcriptome dynamics across and within cell types

Cell types beyond neurons appear to be impacted to some degree by social status. In fact, 13 of the 33 nuclei clusters displayed a significant bias between social states in the relative proportion of nuclei, with five of those cell types being non-neuronal (Fig 1d; Fig S1). One key distinction is that these changes in the proportion of nuclei clusters represent a zero-sum change in the number of cells by social status, so a change more likely reflects a shift in cell transcriptomic state. Additionally, we identified differentially expressed genes in every cluster, ranging from dozens of genes to only a handful in smaller clusters (Fig S2), mirroring the levels of differential expression observed in neuronal nuclei (Fig S3). This aligns with previous work on the role of non-neuronal cell types in regulating social behavior (8). Oligodendrocyte gene expression appears to be particularly sensitive to environmental stimuli, such as previously shown with fasting in mouse hindbrain (52) and hypothalamus (53). There has also been extensive work studying the role of oligodendrocytes in response to social isolation across various brain regions, although less so in the hypothalamus (for review: Tomlinson et al., 2016). This remains an exciting line for future research into how non-neuronal cell types regulate social behavior, especially in brain regions such as the hypothalamus in which much of the work has traditionally focused on peptidergic neurons.

We focused our analyses primarily on the gene networks occurring within neurons due to their direct functional role in regulating brain activity and social behavior, and as neurons are known to exhibit a relatively high degree of transcriptomic variability and subpopulation structure (20). Using a gene co-expression network analysis approach, hdWGCNA (55), we were able to characterize gene networks that go beyond neuronal cell-types or statistical clusters (Fig S6). One assumption that is commonly made about gene networks derived from WGCNA-type approaches in bulk RNA-seq data is that these gene modules reflect cellular-level transcriptomes, hence why those genes are correlated in the first place. As such, we predicted that there would be more modules within a specific neuron sub-type with only a few broader modules representing gene pathways involved in ubiquitous cell function, and therefore, unlikely to be related to social dominance. In fact, and in accordance with our first hypothesis, we found that most modules are highly expressed within specific neuron sub-types (Fig 2c; Fig 6a). This implies that the gene co-expression modules we identified were stratified according to major cell-types, rather than tied into cell state, function, or hypothalamic subregional identity. Additionally, we found that MEG expression differences between dominant and subordinate males depended on whether the analysis encompassed all neuronal nuclei or specific neuronal clusters (Fig 2d; Fig S6b). We interpret these findings such that social status is being actively maintained in both subordinate and dominant individuals through distinct gene networks in separate neuron subtypes. Together, these results indicate that some gene co-expression modules represent specific neuron populations that are highly responsive to social status and/or the social environment.

The gene co-expression modules we uncovered represented a wide range of functions based on enriched GO terms (Dataset S3; Fig S6c; Fig S6d), although functions related to interactions between cells and neuron development were highly prevalent. A key difference in neurons regulating social status may therefore be, at least in part, cell adhesion processes, which requires an understanding of where those cells exist in their neuroanatomical context. Spatial transcriptomics at cellular resolution could provide future insights into relating neuron cellular-level transcriptome dynamics with neuron spatial distribution and neural wiring.

### Arginine vasopressin neuronal subpopulations in relation to social dominance

Across vertebrates, the vasopressin system has long been studied for its role in regulating social behavior (56). We found evidence that there are multiple subpopulations of *avp+* neurons based on the cellular-level transcriptome (Fig 3a). This is consistent with morphological and spatial data identifying multiple subpopulations of *avp+* neurons (18, 49, 57, 58), although we discovered seven distinct molecular cell types of *avp+* neurons, indicating that untangling the relationship within these subpopulations will require further study. Importantly, these results underscore the necessity of integrating multiple lines of evidence across cellular phenotypes from morphology, activity, connectivity, to transcriptomics in determining what is a (functionally distinct) cell type (59).

Because *avp* expression was highest in *avp+* neuron cluster 5 (Fig 3b), this cluster likely contains mostly parvocellular neurons, which are known to have the highest *avp* expression levels (48). Using the HypoMap (21) database as a reference for neuronal nuclei (Fig S4; Fig S5), we correctly predicted the spatial assignment of *avp+* neuron cluster 5 to within the POA (Fig 3c; Fig 3d). This spatial location approach is particularly interesting since in the HypoMap dataset (21) there is a single cluster (C66-22: Caprin2.GLU-6) from the paraventricular nucleus (PVN) that shows the highest amount of gene expression for both avp and oxt (Fig S7a) indicating these are likely be similar to *avp+* subpopulations previously identified in putative teleost homolog (49, 51). We found that *avp+* neuron cluster 4 is majority from neuron cluster 3, which in turn is the best match for HypoMap cluster C66-22 from the PVN. Additionally, *avp+* neuron cluster 6 is majority from neuron cluster 4, and likely represent cells from lateral tuberal nucleus previously described in both *A. burtoni* and midshipman (48, 51). Meanwhile, *avp+* neuron cluster 0 and 4 was spatially assigned to the anterior hypothalamic zone which included the mammalian paraventricular nucleus, which is the putative homolog of the parvocellular cells in the teleost POA (18, 58). While by no means a comprehensive assessment, this reinforces that the statistical clusters found in single-cell data can indeed reflect the cell types found through other approaches and could be expanded towards the study of other candidate genes and pathways.

Previous research in *A. burtoni* demonstrated that the three main morphological subpopulations of AVP*+* neurons – parvo-, magno-, and giganto-cellular – have distinct and social status-dependent *avp* expression patterns (48, 49). Importantly, these status-dependent expression levels are masked when looking at bulk expression of the POA or hypothalamus (48, 50, 60). Likewise, these AVP*+* subpopulations have been shown to differ by social status in other species, such as daffodil cichlids (61). Additionally, *avp*, and not *oxt*, has been shown to have a status dependent role in male social dominance across specific regions of the POA in other species such as the weakly electric fish (62). Surprisingly, we did not find any neuronal *avp+* subpopulations that showed differences in *avp* expression level between dominant and subordinate males (Fig 3b). We did, however, find evidence that the activity patterns of different *avp+* subpopulations may depend on social status (Fig S7c). Previous work in *A. burtoni* demonstrated higher *egr-1* expression following aggressive behavior in *avp+* magnocellular neurons (63). Therefore, the higher *egr-1* relative expression we found in *avp+* cluster 1 of dominant males could be related to the increased aggression seen in these males and potentially indicates that these *avp+* neurons are in fact *avp+* magnocellular neurons. Similarly, we found that *avp+* neuron cluster 5 had higher relative expression of two other IEGs, *npas4* and *fosb*, in subordinate males. This subordinate male activity bias, combined with the higher levels of *avp* expression, further supports *avp+* neuron cluster 5 as a likely candidate for parvocellular *avp+* neurons, which have previously been shown in *A. burtoni* to have higher *avp* expression in subordinate males (48). In fact, *avp+* neuron cluster 5 is mainly comprised of nuclei from neuron cluster 11 (Fig 3c), which also showed the highest IEG expression of any neuron sub-type (Fig S3b). While our study was not designed to test neural activity patterns directly, these differences likely reflect ongoing differences in neuronal activity related to the vastly divergent behavioral repertories associated with social status.

Immunohistochemical studies of AVP and OXT peptidergic neurons have long suggested that these two nonapeptides show little to no co-colocalization (64). In contrast, our study found remarkable co-expression of both *avp* and *oxt* preprohormone mRNA across cell types (Fig S7b). In *A. burtoni* the OXT system has not been studied to the same degree as AVP (65, 66) (but see: (67)), yet similar to *avp*, past work has found no status difference of *oxt* expression when using data bulk POA data (50). While other studies have also showed that *oxt* is broadly expressed in teleosts (68), peptide studies indicate that only a subset of cells in the POA contain the OXT peptide in fish (69–71). Although our findings are not entirely unprecedented as small populations of neurons co-expressing *avp* and *oxt* have previously been shown in rats, specifically in magnocellular neurons (72–75), and similar co-expression has been identified using snRNA-seq in the POA of mice (21, 76). In mammals, the co-expression of *avp* and *oxt* could be related to the close chromosomal proximity (only 3-11 kb apart), with the intervening region known to regulate magnocellular neuron-specific gene expression patterns (77). Previous work demonstrated the regulatory elements driving this cell-specific expression are conserved from pufferfish to rats, even though in pufferfish there is greater physical separation of these genes (46 kb apart with five intervening genes) and a reversal of orientation (78). Since the *avp* and *oxt* genes represent the preprohormones (see below), it is important to consider that post-transcriptional processing produces the two neurophysin co-peptides that facilitate the neurosecretory granule packaging and transport of *avp* and *oxt* prohormone via axons, thereby protecting these neuropeptides from degradation (26, 79–82). Yet, these neurophysins do not appear to co-localize with their opposite neuropeptide counterpart (83), indicating again that *avp* and *oxt* preprohormones may have co-localized transcription but are less likely to have co-localized expression of the mature peptides. In rodents, post-transcriptional processes appear to prevent the co-expression and co-localization of AVP and OXT peptides (see above). More research is needed to determine whether this functional separation is maintained across vertebrates and how it is reconciled with the prevalent co-expression of *avp* and *oxt* preprohormone mRNAs.

### Uncovering the Transcriptomic Neuropeptidome

Neuropeptides and neurosecretory cells date back to metazoan ancestors (24, 25). Prior to single-cell transcriptomic approaches, cellular-level resolution of neuropeptide expression was limited to one or very few genes, making large-scale co-expression analysis difficult (84). Previous single-cell RNA-seq experiments in mouse hypothalamus demonstrated how 27 neuropeptide genes are limited to a few neuronal subtypes, but co-expression of multiple neuropeptides is common in hypothalamic neurons, with some being broadly expressed across subtypes and others more specific (similar pattern found in neuropeptide receptors) (85). With our approach to demultiplexing into genotypes, we demonstrate the consistency of these co-expression networks across dominant and subordinate genotypes and how neuropeptide co-expression reflects social status (Fig 4).

Importantly, expression of a preprohormone does not necessarily lead to mature peptides or guarantee secretion (86). While trying to understand the transcriptomic neuropeptidome, we must recognize that gene expression levels represent preprohormone mRNA, not mature peptide levels. Starting with the discovery that the POMC gene could produce multiple peptides (87–90), it has since become apparent that preprohormones generally encode two or more peptides, which may or may not all be produced in any given cell type (27, 28, 36). snRNA-seq only allows us to estimate preprocessed nuclear mRNA of preprohormones and thus we cannot know which peptide hormones or combination of peptides are ultimately processed and secreted. For example, previous work has demonstrated in *A. burtoni* a dominant male bias in peptide levels of GnRH1 (47) and SST (32), not cell count. Yet, in the present study, we found for both *GnRH1* and *SST* preprohormone mRNA levels a dominant male bias in the proportion of neurons expressing each neuropeptide (Fig 4b). The mechanisms that maintain and regulate peptide translation therefore add an additional layer of complexity to the neuropeptidome. Our discovery of 15 neuropeptide genes with a significant difference in the number of cells in the hypothalamus of dominant and subordinate males highlights the broad potential for this approach. While some of these genes have been shown to have differential expression in *A. burtoni* bulk tissue (male social status in whole-brain transcriptomes *pomc*, *gal, GnRH1* (60); *SST* (32, 89); female reproductive state in hypothalamus transcriptomes *agrp* and *npy* in females (46)), our single-cell dataset opens up many new avenues to explore the role of those less studied neuropeptides.

While previous research has highlighted how specific neuropeptide genes might change either across tissues or on a case-by-case basis at the cellular-level, what remained less clear was whether neuropeptide genes might collectively respond across cells. Therefore, we sought to integrate the entirety of neuropeptide gene expression and co-expression within and across cells, termed the transcriptomic neuropeptidome. This led to the insight that not only are neuropeptide-levels changing with social status, but that the co-expression of specific pairs of neuropeptides also changes (Fig 4c; Fig 4d; Fig S8). Additionally, our analysis of the transcriptomic neuropeptidome also revealed that dominant males have both a more highly integrated and consistent neuropeptidome relative to subordinate males. This implies that the maintenance of a dominant phenotype requires molecular pathway to be more tightly controlled compared to the more variable subordinate males. In fact, this elevated gene network integration pattern in dominant male *A. burtoni* has previously been demonstrated with a few select neuropeptides (50, 91). The combinatorial nature of these cellular-level neuropeptide gene networks, taken together with this more integrated network, implies that social status reflects more than simply the activation of a few well-studied gene pathways but a complex interplay across the genome.

## CONCLUSIONS

We used single-nucleus RNA-seq in the hypothalamus and POA of A. burtoni to understand the effect that social status has on cellular gene expression. We demonstrate how social status manifests from changes in broader neuronal gene networks to targeted changes between known socially relevant neurons. We assess the entirety of the transcriptomic neuropeptidome to understand the interaction of neuropeptide gene networks with social dominance. These findings provide a rich dataset for future functional work and a conceptual framework for comparative studies on the evolution of the neural mechanisms of social behavior. Insights into the transcriptomic networks that modulate social status, specifically with neuropeptides, aid in our understanding of the complexity inherent in social behavior.

## MATERIALS AND METHODS

### Animals

The animals used in this study came from a laboratory population of Burton’s Mouthbrooder cichlid, *Astatotilapia burtoni*, descended for about 60 generations from a wild-caught stock of 400 individuals (92). All work was done in compliance with the Institutional Animal Care and Use Committee (IACUC) at The University of Texas at Austin.

### Experimental design

We created naturalistic laboratory communities of adult *A. burtoni*, with two males and three females each, to generate three pairs of dominant and subordinate males used to compare snRNA-seq of the hypothalamus and POA by social status (Fig 1a). Detailed experimental methods are described in the Supplemental §1 (Dataset S4).

### Processing of single nucleus data

We obtained 926 million reads total, with ∼460 million reads for the dominant samples pool and ∼466 million reads for the subordinate samples pool. We used the MultiQC package (93) to confirm that all reads had Phred scores above 35 and passed all quality control measures. Read mapping and annotation were implemented using the Cell Ranger 6.1.2 pipeline (94) and the closely-related Nile tilapia, *Oreochromis niloticus*, reference genome (O_niloticus_UMD_NMBU) (95). For the subordinate pool, 74.1% of reads mapped to the genome, and 27,726 total genes were detected. The cell by gene expression matrices for each pooled sample dataset were filtered by selecting genes expressed in a minimum of 3 cells and selecting nuclei with 1000-2000 detected genes (nFeature_RNA). This threshold was determined from the distribution of nuclei to capture the bulk of the nuclei distribution while removing the tails to reduce background and reduce doublets to get the highest quality 10,000-20,000 nuclei per sample, while the metric consistent across sample libraries. Additionally, nuclei were filtered for total molecules (nCount_RNA) less than 6,000 and percentage of mitochondrial genes less than 5%. This resulted in a total of 14,534 and 12,243 single-nucleus transcriptomes for the dominant and subordinate sample pool respectively. For the dominant pool, there were 1,237 median genes per nuclei and 1,776 median UMI counts per nuclei after filtering. For the subordinate samples pool, there were 1,246 median genes per nuclei and 1,785 median UMI counts per nuclei after filtering. Data was then processed with the Seurat integrated analysis pipeline (96) and integrated with SCTransform version 2 (97) with the 3,000 most variable genes, we utilized Clustree (98) to determine an appropriate resolution for clustering by selecting a resolution that yielded the most stable number of clusters (0.8).

### Genotype demultiplexing

Even though our *A. burtoni* laboratory population is not fully outbred (99, 100), we expected there to be sufficient genotypic variation to allow demultiplexing of pooled samples by genotype, thereby recovering biological replication for downstream statistical analyses. The inclusion of greater biological variation with identification of multiple genotypes can further reduce any batch effects, although the main source would likely be due to sequencing and prep which were done at the same time. Genotypic variation in our *A. burtoni* populations allowed for demultiplexing of the pooled samples using genotype of each sample, although genotypes are not tied directly to specific individual animals. We used the souporcell package (101) for this purpose, as it does not require the genotype of each individual in a pooled samples to be known. Briefly, the pipeline performs variant detection on the sequence bam files to count allele support for each nucleus. Next, the nuclei are clustered with a sparse mixture model to assign cells to genotypes. Together, this allows for demultiplexing in the absence of prior genotype information of each sample. Demultiplexing identified three dominant and three subordinate genotypes, with dominant males comprising 4845 ± 414 (mean ± SE) nuclei and subordinate males 4081 ± 110.

### Cell type calling

We successfully recovered all major cell types previously identified in the mammalian hypothalamus (See Supplemental §2). Importantly, this approach allowed us to reduce noise by characterizing nuclei with unknown cell types that did not get assigned to any specific cell category. We compared the proportion of cell subtypes between dominant and subordinate males with a binomial GLM for each cell subtype separately (102) and FDR-corrected to account for comparisons across all the subtypes we tested. To assign putative spatial location to neuron clusters in *A. burtoni*, we developed a method for comparing these clusters to mouse hypothalamus neuron clusters with known or imputed spatial information on the HypoMap database (21) (See Supplemental §3). We took this comparative approach for spatial localization because a recent spatial transcriptomic atlas for another cichlid species (103) is focused on the forebrain and lacks cellular resolution, which makes direct comparison challenging.

### Differential gene expression analysis and weighted gene co-expression network analysis

We further subclustered each cell type, mediating identification of subtypes, using UMAP and Clustree (98) to determine the appropriate number of sub-clusters. To identify DEGs in these clusters according to social status, we used limmatrend (104, 105) in a pseudo-bulk framework, aggregating expression within each cluster for biological replicates while incorporating genotype to increase statistical power. This approach has previously been shown to be robust (105) and has been implemented in the R package Libra (106). First, we filtered lowly expressed genes by requiring greater than 10% detection in at least one cluster in at least one social status, to a total of 6,006 genes prior to running DEG analysis on all clusters. For genes identified as differentially expressed by pseudo-bulk limma (FDR-corrected p < 0.05), we additionally quantified the proportion of cells expressing each gene (defined as RNA counts >2) within each cluster by social status to help disentangle whether DEGs were from changes in expression level within nuclei or changes in the number of nuclei expressing the gene. Concordance in effect direction between these proportion difference estimates and the DEG results aided the biological interpretation. Yet, as social status and library are colinear in our data, differential expression results are considered exploratory.

Nuclei identified as GABAergic “inhibitory” (GABA) and glutamatergic “excitatory” (GLU) were merged into the broader neuron category. Due to the higher degree of complexity of neurons, indicated by the presence of multiple sub-clusters, these neuron nuclei were subsequently subset out of the dataset and re-clustered with UMAP into various sub-populations of neurons. The proportion of neuron subtypes was compared between dominant and subordinate males with a binomial GLM for each cell subtype separately (102) and FDR corrected to account for comparisons across all subtypes tested.

Next, we performed a Weighted Gene Co-expression Network Analysis (WGCNA) approach adapted for single-nucleus data, hdWGCNA (55), to identify gene co-expression modules for the 3,000 most variable genes. Compared to traditional WGCNA (107), hdWGCNA was developed specifically for use with single-cell and spatial transcriptomics data by generating metacells from selected pools of nuclei to reduce background noise of typically noisy single-nucleus transcriptomes. These metacells represent localized groups of cells distinct from seurat clustering, using a K-nearest neighbors (KNN) algorithm, producing an expression matrix that has less sparsity and, therefore, more amenable to correlation network approaches. Metacells were pooled by genotype (see above for details) and with a minimum cell count parameter of 50. We then examined whether and how modules differed by social status using differentially expressed module eigengenes (MEGs) with Wilcoxon rank-sum from hdWGCNA (55). To determine whether modules were specific to certain neuron clusters, MEGs were averaged across neuronal clusters independent of social status. Modules were considered variable across neuron clusters if the standard deviation of the neuron cluster mean MEGs divided by the mean MEG across clusters was greater than 1. Cluster specificity was defined by z-scored cluster means greater than 2. Modules that were both variable and specific across neuron clusters were used for differential testing by social status. To assess MEG expression differences within specific neuron clusters, ME values were pseudo-bulked by averaging across nuclei within each cluster and genotype. Differential MEG expression by social status within specific module-neuron cluster pairs was assessed using linear models applied to mean pseudo-bulk expression values by genotype weighted by relative nuclei counts and adjusted with FDR.

To determine whether modules were functionally enriched for specific gene ontology (GO) terms, we used AnnotationForge to generate a GO database for Nile Tilapia (108), calculated GO enrichment with enrichGO from the package clusterprofiler (109), with FDR corrected adjusted p-value cutoff of 0.15, minGSSize = 10, and maxGSSize = 500, and graphed our results the package enrichplot (110).

To assess neuronal activity, we quantified the proportion of neurons expressing immediate early genes (IEGs). First, we calculated the percentage of nuclei expressing IEGs (*egr1, npas4, fosb, fosab*) as well as a composite IEG score for each genotype across neuron clusters filtering out clusters with less than 5% of nuclei IEG+ in at least one condition. In each cluster, differences in the proportions of IEG expressing nuclei were assessed across genotypes using a two-sample t-test with FDR correction. Significance was defined as either FDR p < 0.15 or 95% confidence interval excluding zero.

### Neuropeptide preprohormone co-expression networks

To infer the structure of the neuropeptidome, the entirety of neuropeptide genes expression within a cell, we collated known “neuropeptide genes” (encoding pre-prohormone mRNAs) from several databases (33–35) and via Ensembl (111), resulting in 127 genes (Dataset S5). To assess the transcriptomic neuropeptidome we generated co-expression network of just neuropeptide-encoding genes for each genotype. We used a dynamic threshold to determine presence/absence of a neuropeptide gene in a cell to better capture both genes expressed rarely and genes expressed broadly at low levels across many nuclei. Therefore, rarely expressed genes (<10% of neuronal nuclei) were considered present if they were expressed, whereas for highly expressed genes (>30% of neuronal nuclei) presence was only assigned to nuclei in the top 15% of expressing nuclei, while moderately expressed genes (between 10% and 30%) presence was only assigned to nuclei in the top 30% of expressing nuclei.

We used an Anderson-Darling k-Sample Test to determine whether the number of neuronal nuclei that expressed different numbers of neuropeptides came from the same distribution within and between social states (112). To test whether the distributions of neuropeptide genes per neuronal nuclei differed across and within social status we used Poisson regression. The proportion of neuronal nuclei expressing each neuropeptide gene was compared between dominant and subordinate males with a binomial GLM for each neuropeptide separately (102) and adjusted for multiple comparisons within subtypes with a multivariate t distribution with the R package ‘glht’ and adjusted again with FDR to account for comparisons across all subtypes tested.

To identify gene pairs that differed by social status, for each edge, representing a neuropeptide gene pair, an FDR corrected t-test was used to determine if the specific co-expression pair was dominant and subordinate biased. Global and local clustering coefficients were calculated with transitivity in igraph (113).

To compare transcriptomic neuropeptidome similarity by social status, we first filtered networks to neuropeptides expressed in at least 25 neuronal nuclei across all samples. Using the same co-presence networks normalized by cell number for each genotype, we calculated Pearson correlation among all pairwise genotypes. We then calculated mean and standard deviation for pairs within and across social states, using a wilcox test to compare within and across status. Additionally, we compared networks by social status using the permutation approach PERMANOVA (N=719) with the function adoni2 and betadisper from the package vegan (114).

## DATA AVAILABILITY STATEMENT

The filtered deduplicated reads have been deposited to the NCBI Short Read Archive (SRA), bioproject PRJNA1310498. The code used in this paper together with documentation can be accessed via Zenodo (115)

## ACKNOWLEDGEMENTS

We thank Dr. Nihal A. Salem for guidance and the members of the Hofmann Lab for discussion and Holly Stevenson for technical assistance. We also thank Dr. Zackary V. Johnson and Dr. Tessa K. Solomon-Lane for helpful suggestions and feedback on earlier version of the manuscript. Sequencing was performed by the Genomic Sequencing and Analysis Facility at UT Austin, Center for Biomedical Research Support (RRID# SCR_021713). Computational analyses were performed on the Biomedical Research Computing Facility at UT Austin, Center for Biomedical Research Support (RRID#: SCR_021979).

