## Supplemental for "Social Dominance Reorganizes the Transcriptomic Neuropeptidome in a Highly Social Cichlid Fish"

**This PDF file includes:**

Legends for Datasets S1 to S5

Supplemental Methods §1 to §3

Supplemental References

Figures S1 to S8

### Supplemental Table Legends

**Dataset S1 (separate file).** Results of binomial GLM comparing proportion of nuclei across social status for overall nuclei clusters (Seurat_clusters), neuron clusters (Neuron_clusters), and for individual neuropeptide genes (Neuropeptide_genes). Each datasheet includes the contrast, estimate, standard error, z value, p-value, FDR adjusted p-value. For both the overall nuclei clusters and neuron clusters, also included are the upper and lower 95% confidence intervals.

**Dataset S2 (separate file).** Pseudobulk limma DEG results for broad nuclei clusters (all.deg.results) and neuronal nuclei clusters (neuron.deg.results) with cluster name (cell_type), gene, average log fold change (avg_logFC), percent of subordinate nuclei expressing gene (sub_burtoni_snseq.pct), percent of dominant nuclei expressing gene (dom_burtoni_snseq.pct), average gene expression level of subordinate nuclei (sub_burtoni_snseq.exp), average gene expression level of dominant nuclei (dom_burtoni_snseq.exp), p-value (p_val), FDR adjusted p-value (p_val_adj), and differential expression family (de_family), method (de_method), and type (de_type).

**Dataset S3 (separate file).** GO enrichment analysis results for hdWGCNA modules for neuronal nuclei with module name, GO ID, GO description, ratio of annotated genes in a term (GeneRatio), ratio of all genes in annotated term (BgRatio), p-value, adjusted p-value, q-value, list of gene names in GO term (geneID), and number of genes (Count).

**Dataset S4 (separate file).** Sample metadata including information on size (standard length and mass), dissection, gonads, and testosterone.

**Dataset S5 (separate file).** Gene lists for known cell type marker genes from mammalian literature (Mammalian_literature_markers), Nile Tilapia orthologs assigned to HypoMap orthologs (HypoMap_tilapia_markers), and neuropeptide gene list (Neuropeptide_gene_list) collated from multiple mammalian databases with assigned Nile Tilapia orthologs.

### **Supplemental Methods**

§1. Detailed experimental design

*Sample collection*

We created six naturalistic communities of adult *A. burtoni* males and females within three 100-L aquaria, with each aquarium subdivided by a transparent, perforated divider to facilitate visual and chemical communication between neighboring communities. Each community consisted of one large male (57-61 mm standard length) and one male that was 5-10 mm smaller (45-53 mm standard length), as well as three females (10-15 mm smaller than the large male). A terracotta pot served as territorial shelter and artificial aquarium plants provided a refuge for the subordinate male and females. We performed live global scoring of social status and coloration of every male in each community every other day for 16 days. We also recorded reproduction from the presence of any mouthbrooding females. After three weeks, we collected all males between 10:00h and 12:00h, measured their body mass and standard length, and collected blood for subsequent hormone assays. Males were then euthanized via rapid cervical dissection and brains were dissected within three minutes, embedded in OCT compound (Sakura Finetek, USA), immediately frozen on dry ice, and stored at -80°C until further processing. Gonads were dissected and weighed to obtain the gonadosomatic index, a proxy measure of reproductive state, by dividing gonad mass by body mass. Blood samples were spun in a table centrifuge for five min at 3000 rpm to separate the plasma, which was stored at -80°C until further processing.

*Social status assessment*

From the six communities, we selected 3 pairs of dominant and subordinate males that displayed a stable social phenotype over the 16-day observation period. Dominance was assessed by the display of traits typically associated with social dominance in *A. burtoni* such as: aggressive and courtship behavior, conspicuous yellow or blue body coloration, and the presence of an eye bar (1, 2). We only selected subordinate males that were never observed to show a dominance trait. Since dominant males perform reproductive behavior and suppress that same behavior in subordinates, we required at least one occurrence of a brooding female in the community, indicating that the dominant male had mated successfully at least once. As community setup included a bias in size to help ensure a more stable dominance relationship, dominant males were on average larger by 23% in standard length ($t(2) = 5.05$, $p = 0.037$) and 79% in mass ($t(2) = 23.21$, $p = 0.002$) compared to subordinate males. We additionally used hormonal differences to help distinguish status in each pair by higher circulating testosterone levels in dominant males ($M = 94$ ng/mL, $SD = 52$) than subordinate males ($M = 9.6$ ng/mL, $SD = 14$) ($t(2) = 2.72$, $p = 0.098$), although we found no difference in GSI mass ($t(2) = 0.896$, $p = 0.465$).

*Hormone assay*

Circulating testosterone levels were assayed in duplicate from blood plasma by ELISA (Cayman Chemical, USA) on a single assay plate following manufacturer's instructions (within-plate coefficient of variation was 12%). Note that in contrast to many other teleost fishes, testosterone is the main androgen in haplochromine cichlids (3, 4).

*Tissue collection*

The POA and hypothalamus were micro-dissected following the protocol described in (5). Briefly, brains were sectioned into 300 µm slices on a cryostat microtome onto positively charged Superfrost Plus microscope slides (Thermo Fisher Scientific, USA), which were then placed on a metal block chilled on dry ice for 10 min, or until temperature reached approximately -4 to -10 °C, to allow the collection of brain tissue without fracturing sections. Using a 0.75 mm tissue punch tool (Electron Microscopy Sciences, USA), the entirety of the POA and hypothalamus was collected together into a 1.5 ml reaction tube (16-18 punches per subordinate and 20-26 punches per dominant) and stored at -80°C until further processing.

*Nuclei isolation*

Neural tissue is challenging to dissociate into nuclei due to the heterogenous nature of the tissue and unique interconnected structure of neurons. Fluorescence-activated cell sorting (FACS) is a common method to isolate and purify single cells, for multiple parameters and cell-types, but requires a starting volume larger than micro-dissected brain tissue punches can provide (6). It has been shown that the transcriptomes of single cells and single nuclei are highly correlated (7). We therefore opted to isolate single nuclei from frozen tissue instead.

It should also be noted that statistical power of single-cell sequencing is largely determined by the number of cells (nuclei) and sequencing complexity, rather than sequencing depth (6, 8, 9). Pooling samples increases statistical power by increasing the number of nuclei while still capturing the biological variation between samples. Pilot experiments indicated that pooling the entire POA and hypothalamus together for three males would be required to reach a minimum number of 10,000 cells (or nuclei) per sample, as recommended for the 10X Genomics Chromium platform (10x Genomics, USA). We therefore selected one dominant-subordinate pair from each of our three separate aquaria and combined them into two separate pools for dominant and subordinate males, respectively (1 pool per group, n=3 animals per pool).

We adapted a density gradient protocol for nuclei isolation (10). Briefly, all tissue samples were frozen or on ice throughout the protocol to prevent any additional transcriptional activity. First, the tissue was mechanically homogenized to release the nuclei by adding 1.5 ml of chilled Nuclei EZ Lysis Buffer (Sigma-Aldrich, USA) with 0.2 U/µl Protector RNase Inhibitor (Roche, USA) to the tissue in a 2 mL tube and then gently homogenized 20 times in a chilled 7 ml Dounce homogenizer with pestle "b." Nuclei were then filtered through a 35 µm cell strainer cap from Falcon Round-Bottom Polystyrene Test Tubes with Cell Strainer Snap Cap, 5mL (Corning, USA) into a 1.5 mL tube. Next, the nuclei were centrifuged at 900 g for 5 min at 4°C before the supernatant was removed and the nuclei pellet resuspended in 750 µl of 25% Optiprep (Sigma-Aldrich, USA) in nuclei resuspension buffer of 1x PBS with 2% BSA and 0.2 U/µl Protector RNase Inhibitor (Roche, USA). Resuspended nuclei were then overlaid on 750 µl of 29% Optiprep in nuclei resuspension buffer before centrifuging at 13,000 g for 30 min at 4°C. Afterwards, the supernatant was carefully removed, and the pellet quickly resuspended with 20 µl nuclei resuspension buffer and transferred to a new 0.5 ml tube with 50 µl of nuclei resuspension buffer added. Nuclei were then filtered through a 35 µm cell strainer cap (Corning, USA). Finally, the concentration of nuclei was quantified by using two 5 µl aliquots with 5 µl resuspension buffer with 10 ug/mL DAPI on the Countess 3 Automated Cell Counter (Invitrogen, USA) to aim for a concentration between 700-1200 nuclei/µl. The samples were then submitted to UT Austin's Genomic Sequencing & Analysis Facility for quality control and library construction using the 10X Genomics Chromium Next GEM Single Cell 3' reagent kit v3.1 as per manufacturer's instructions. The libraries were then sequenced on the Illumina NovaSeq S1platform.

§2. Cell type annotation

The lack of standardized marker genes as well as the possible presence of multiple gene paralogs complicate cell type identification in non-traditional model systems. A list of known cell type marker genes was obtained from the mammalian literature and another marker list was obtained from HypoMap (C7 markers) (11), a murine hypothalamus single-cell expression atlas from an 18 study meta-analysis (Dataset S5). Recent work has demonstrated the deep homology of cell type expression between teleosts and other vertebrate lineages in the forebrain, lending credence to this cross-species comparison (12). We determined Nile Tilapia orthologs of these genes with ensembl. We tested a few different methods to identify distinct cell type classes including: Seurat's Module score (13), scsorter (14), and sctype (15). To best characterize nuclei with unknown cell types, and to robustly identify cell types, we used the combined set of cell marker gene orthologs and performed cell-type identification with sctype, an ultra-fast and automated cluster-based method. Importantly, sctype default assigns a cell-type to a cluster if the top positive scaled cell-type score for the entire cluster is greater than 25% the number of cells in the cluster. Due to differences in the number of orthologous marker genes for each cell-type marker from HypoMap, these scores can be artificially low for certain cell types and particularly impacts neurons. To account for this, if a cluster was assigned unknown but the top cell-type score was positive for neuron-type that matched the marker gene pattern and the cell type tree, the cell was assigned the cell-type neuron-type. Only three nuclei clusters (clusters 2, 6, and 7) were impacted by this lower threshold. Otherwise, clusters were assigned as unknown.

§3. Spatial location prediction

The hypothalamus represents a brain region comprised of multiple, distinct, spatially distributed sub-populations with known functional differences. To assign putative spatial location to neuron clusters in *A. burtoni*, we developed a method for comparing these clusters to mouse hypothalamus neuron clusters with known or imputed spatial information. Predicting cluster specific spatial location relied on the HypoMap database (11), which uses a combination of the *in situ* hybridization (ISH) expression values from the Allen Brain Atlas (16), and described region of origin from the original dataset. First, we selected neuron cluster marker genes compared to all other neuron clusters with FindAllMarkers (min.pct = 0.1) in Seurat (13). A specificity score, as described in Steuernagel et al, 2022 (11), was calculated for each marker gene. Briefly, specificity was calculated as the average expression fold-change multiplied by the ratio of the proportion of expressing nuclei within the cluster to that across all neurons for each marker gene of a given cluster. Additionally, HypoMap cluster marker genes for neurons in clusters 1-50 of C66 were mapped to Nile Tilapia orthologs with Biomart (17). These HypoMap cluster marker genes include a specificity score as well. Due to limited overlap of orthologous genes between the marker gene sets for neuron clusters and HypoMap C66 clusters (Fig S4a), we filtered out clusters with less than 10 cichlid orthologs assigned as marker genes. We then used pairwise Spearman correlation between marker gene specificity score for the neuron clusters and the HypoMap cluster using the shared orthologs. To assess significance of these correlations, we created a null distribution for each neuron cluster with a permutation approach (n = 1,000), by randomly shuffling gene specificity scores and then recalculating the correlations with HypoMap clusters. From this, observed correlations could be converted into permutation z-scores, that we then removed low-confidence values by setting all z-scores below 1 to 0. To assign brain regions, we took advantage of the fact that HypoMap clusters are comprised of cells from known hypothalamic brain regions (Fig S4b) and used the proportion of cells from each brain region to annotate each HypoMap cluster. For each neuron cluster and HypoMap cluster pairing, z-score values were weighted by the proportion of cells from known brain regions and aggregated across regions. Therefore, this weighted z-score reflects the likelihood that a neuron cluster is comprised of a certain brain region as annotated by HypoMap. To further enhance annotation, regions were collapsed into broad hypothalamic spatial zones (Preoptic, Anterior, Ventral, Dorsal, Lateral, Posterior) (Fig S4b), and weighted z-scores were summed within each spatial zone for each neuron cluster.

### Supplemental Figures

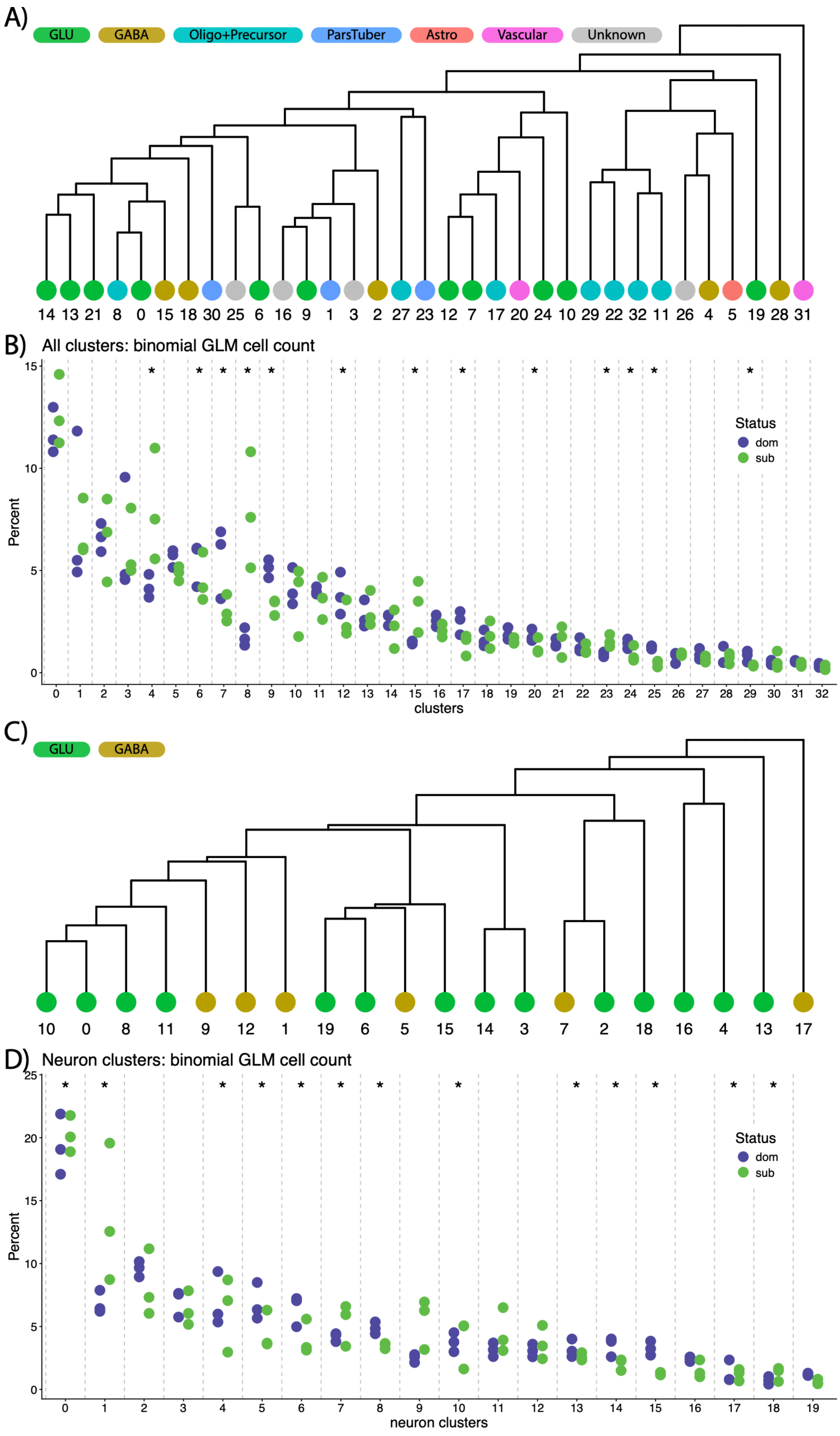
A)
GLU
GABA
Oligo+Precursor
ParsTuber
Astro
Vascular
Unknown
14 13 21 8 0 15 18 30 25 6 16 9 1 3 2 27 23 12 7 17 20 24 10 29 22 32 11 26 4 5 19 28 31
B) All clusters: binomial GLM cell count
Percent
clusters
Status
dom
sub
C)
GLU
GABA
10 0 8 11 9 12 1 19 6 5 15 14 3 7 2 18 16 4 13 17
D) Neuron clusters: binomial GLM cell count
Percent
neuron clusters
Status
dom
sub

**Supplemental Figure 1.** Cell-type annotation and genotype percentages across clusters. A) Hierarchical dendrogram of all cell type clusters from transcriptome data labeled with cluster annotation. B) Percentage of total nuclei assigned to each individual genotype for dominant (purple) and subordinate (green) males in each cluster. C) Hierarchical dendrogram of neuron cell type clusters from transcriptome data labeled with neuron cluster annotation. D) Percentage of total nuclei assigned to each individual genotype for dominant (purple) and subordinate (green) males in each neuron cluster. Clusters with significant difference ($p$-value $< 0.01$) in social status from binomial GLM denoted with *.

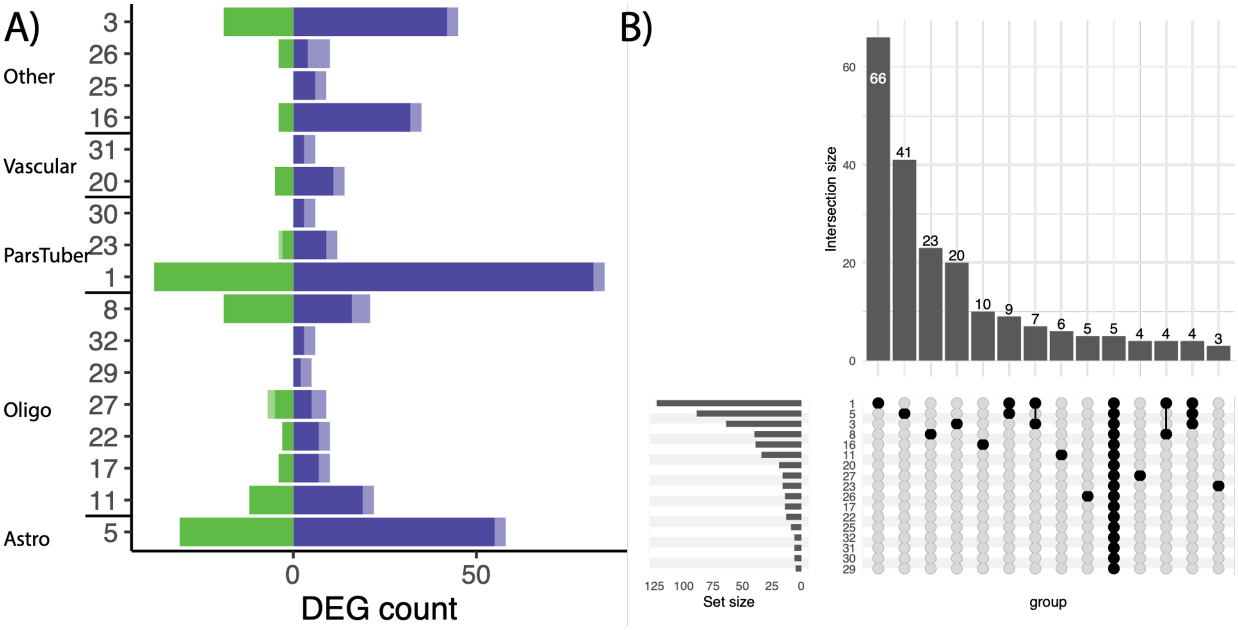


**Supplemental Figure 2.** Non-neuronal DEG. A) Number of significant (FDR < 0.05) DEGs (x-axis) in each non-neuronal cluster (y-axis) that are higher in dominant (purple) or subordinate (green).that are associated with a difference in the percent of nuclei expressing greater than 20% across social status for that gene in a cluster are designated with a lighter color on the end of each bar. B) Upset plot showing the overlap of DEGs across non-neuronal clusters.

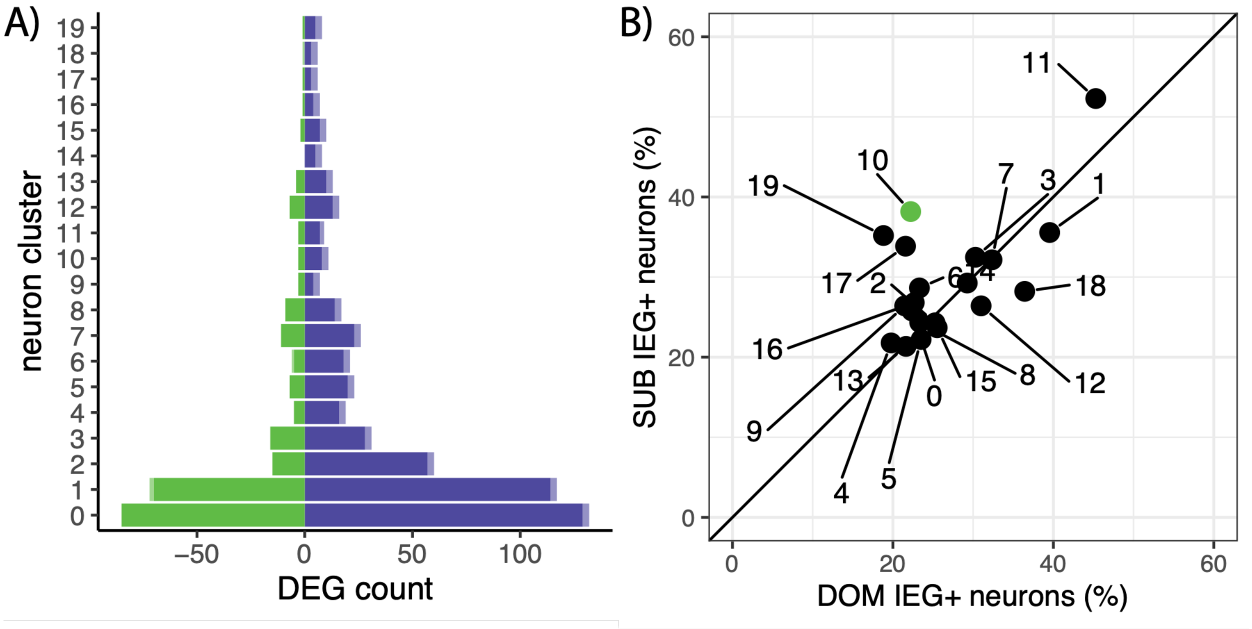


**Supplemental Figure 3.** Neuron DEG and IEG+. A) Number of significant (FDR < 0.05) DEGs (x-axis) in each neuron cluster (y-axis) that are higher in dominant (purple) or subordinate (green). DEGs that are associated with a difference in the percent of nuclei expressing greater than 20% across social status for that gene in a cluster are designated with a lighter color on the end of each bar. B) Proportion of nuclei that are IEG+ for each neuron cluster. Color indicates significant difference (FDR < 0.1) in proportion of IEG+ nuclei across social status (green = subordinate, purple = dominant).

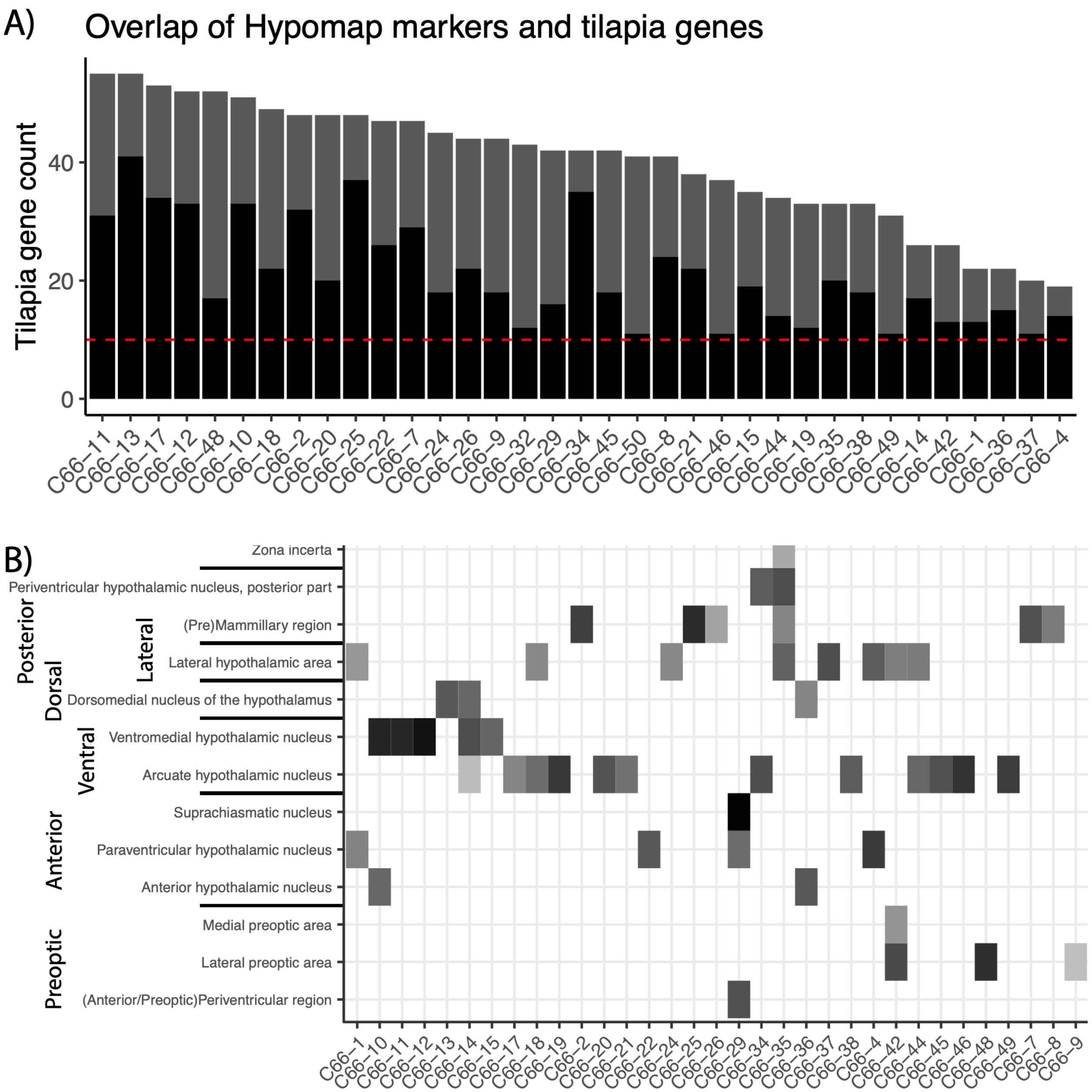


**Supplemental Figure 4.** Hypomap spatial location. A) The number of marker gene orthologs in tilapia from mouse (grey bar) for each hypomap clusters (x-axis) compared to the number of tilapia marker gene orthologs present in the *A. burtoni* single-nuclei data (black bar). Red line indicates cutoff threshold of 10 to filter out hypomap clusters. B) Each hypomap cluster contains nuclei identified as originating from specific brain regions. Box color indicates number of nuclei from a given hypomap cluster (x-axis) assigned to a given brain region (y-axis). Horizontal black bars on the y-axis indicate which regions were grouped together to form broader hypothalamic spatial zones.

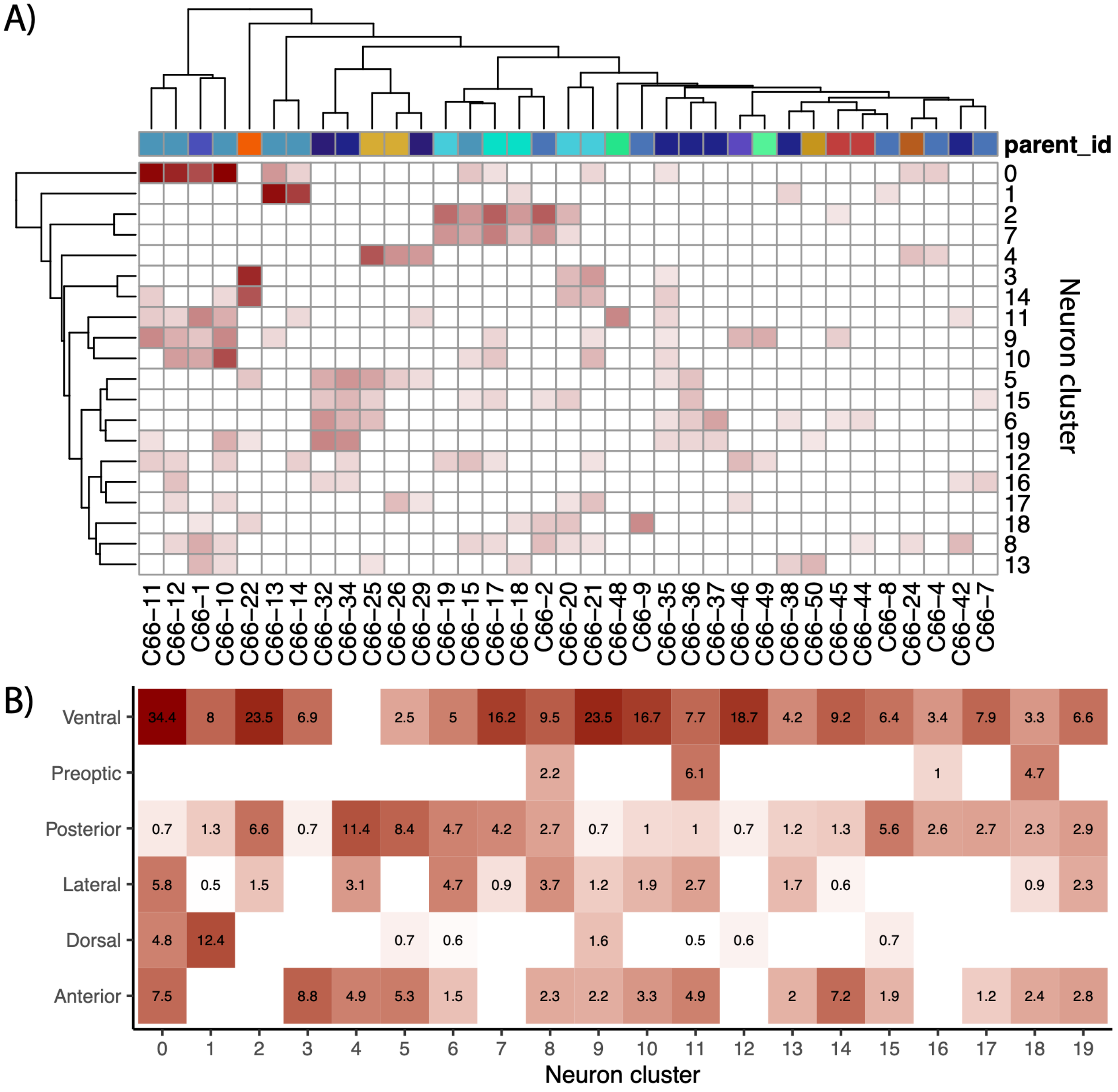


**Supplemental Figure 5.** Neuron cluster brain region identification with HypoMap. A) Heatmap of weighted permutation z-scores pairings between neuron cluster (rows) and HypoMap clusters (columns). Additionally, HypoMap clusters are labeled by the hierarchical clusters each of the C66 clusters is assigned from (top row colors; parent_id). B) Calculated likely brain region scores for each neuron clusters when comparing specificity score of overlapping genes to HypoMap clusters and accounting for proportion of cells assigned to a certain brain region.

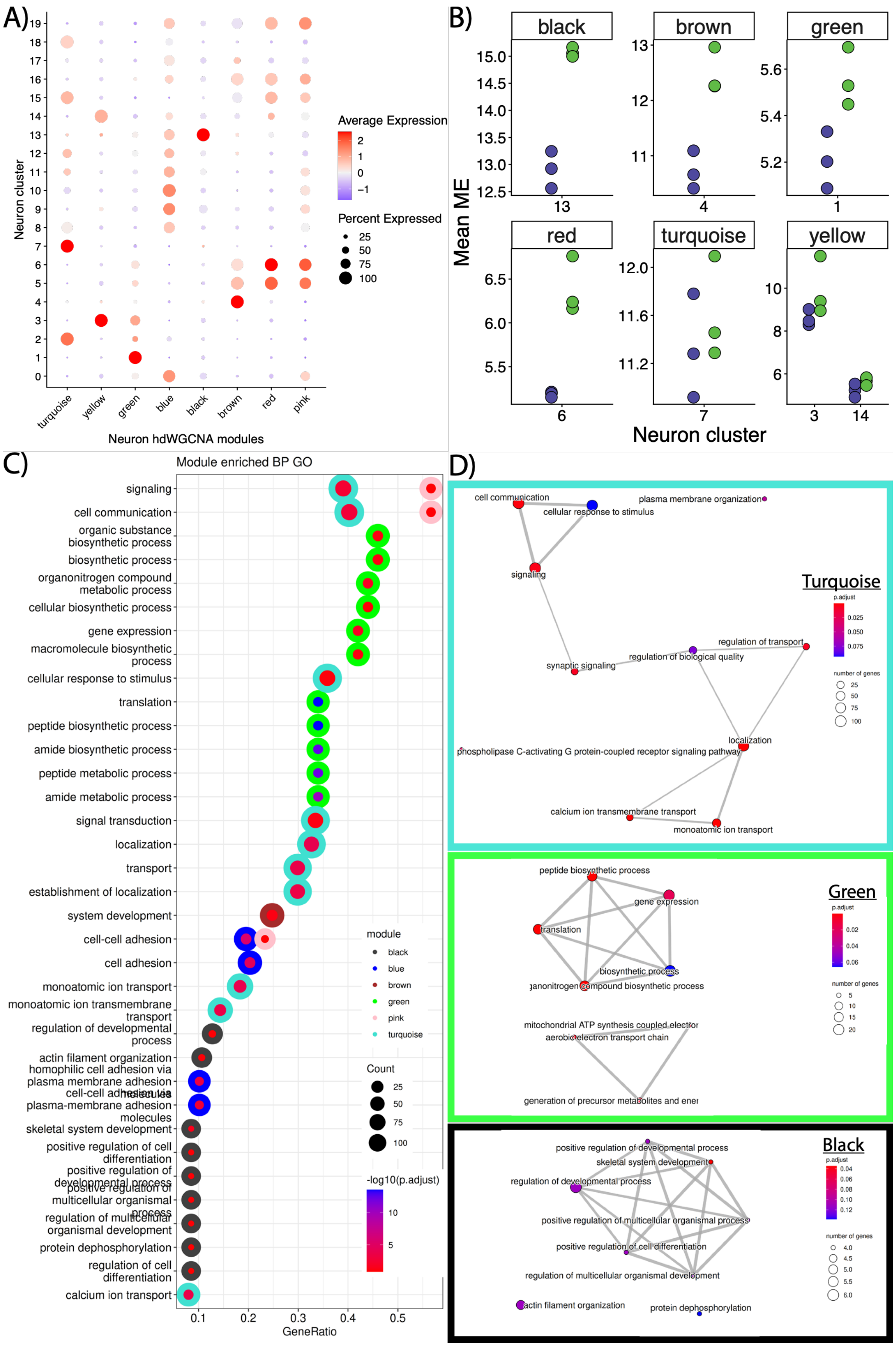

A)
B)
C)
D)
Average Expression
Percent Expressed
Neuron cluster
Neuron hdWGCNA modules
turquoise
yellow
green
blue
black
brown
red
pink
Mean ME
Neuron cluster
Module enriched BP GO
GeneRatio
module
Count
-log10(p.adjust)
signaling
cell communication
organic substance biosynthetic process
biosynthetic process
organonitrogen compound metabolic process
cellular biosynthetic process
gene expression
macromolecule biosynthetic process
cellular response to stimulus
translation
peptide biosynthetic process
amide biosynthetic process
peptide metabolic process
amide metabolic process
signal transduction
localization
transport
establishment of localization
system development
cell-cell adhesion
cell adhesion
monoatomic ion transport
monoatomic ion transmembrane transport
regulation of developmental process
actin filament organization
homophilic cell adhesion via plasma membrane adhesion
cell-cell adhesion via plasma-membrane adhesion molecules
skeletal system development
positive regulation of cell differentiation
positive regulation of developmental process
positive regulation of multicellular organismal process
regulation of multicellular organismal development
protein dephosphorylation
regulation of cell differentiation
calcium ion transport
Turquoise
Green
Black
p.adjust
number of genes
cell communication
plasma membrane organization
cellular response to stimulus
synaptic signaling
regulation of biological quality
regulation of transport
phospholipase C-activating G protein-coupled receptor signaling pathway
calcium ion transmembrane transport
monoatomic ion transport
peptide biosynthetic process
biosynthetic process
mitochondrial ATP synthesis coupled electron
aerobic electron transport chain
generation of precursor metabolites and ener
positive regulation of developmental process
skeletal system development
regulation of developmental process
positive regulation of multicellular organismal process
positive regulation of cell differentiation
regulation of multicellular organismal development
actin filament organization
protein dephosphorylation

**Supplemental Figure 6.** hdWGCNA module analysis. A) Dotplot depicting module eigengene average expression (color) and percent of nuclei expressing (size) for each neuron hdWGCNA module (x-axis) across the neuron clusters (y-axis). B) Comparison between dominant (purple) and subordinate (green) males average module eigengene expression (y-axis) across individual genotypes for neuron cluster-specific modules. C) GO enrichment analysis to identify significantly (FDR corrected p-value < 0.15) enriched GO terms for the GO category biological processes in each of the hdWGCNA neuron modules. Dotplot indicate GO terms (y-axis), GO term gene ratio (x-axis), number of genes for that GO term (inside dot size), the FDR corrected p-value (inside dot color), and the module being tested (outside ring color). For each module, GO terms were selected to include the GO terms with the highest gene counts and most significant by adjusted p-value D) Network diagram of semantic similarity for all significantly enriched GO terms in a module (top – turquoise; middle – green; bottom - red), with closely-related GO terms collapsed together.

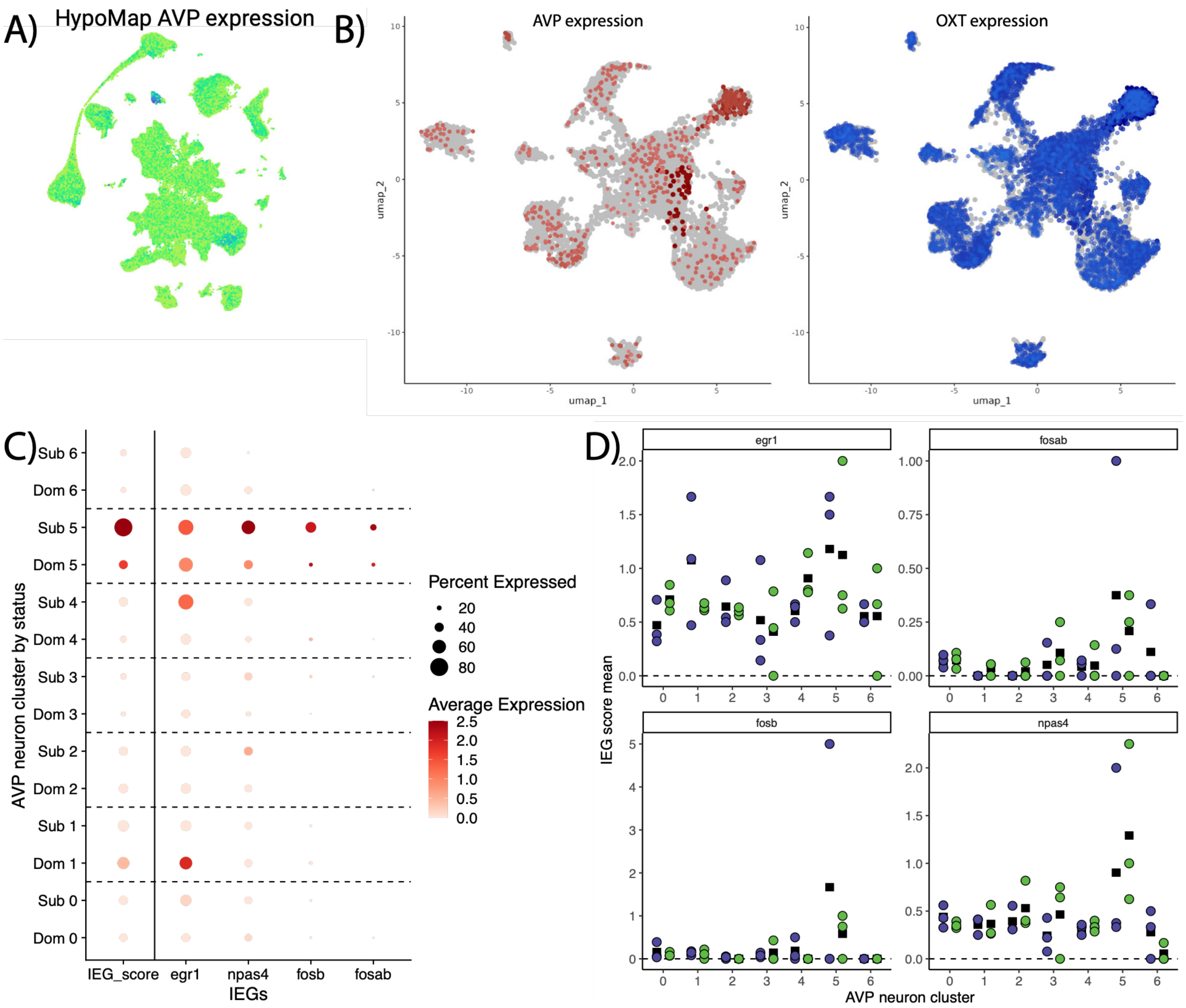


**Supplemental Figure 7.** AVP neurons gene expression. A) HypoMap scvi projection of AVP gene expression across all nuclei. B) Neuron UMAP projection with *avp* (left) and *oxt* (right) expressing neurons shaded by the average relative expression for each AVP neuron cluster. C) Dotplot of immediate early gene (x-axis) expression levels across AVP neuron clusters by social status (y-axis). D) Average immediate early gene expression (y-axis) of AVP neuron clusters (x-axis) by individual across social status (green = subordinate, purple = dominate).

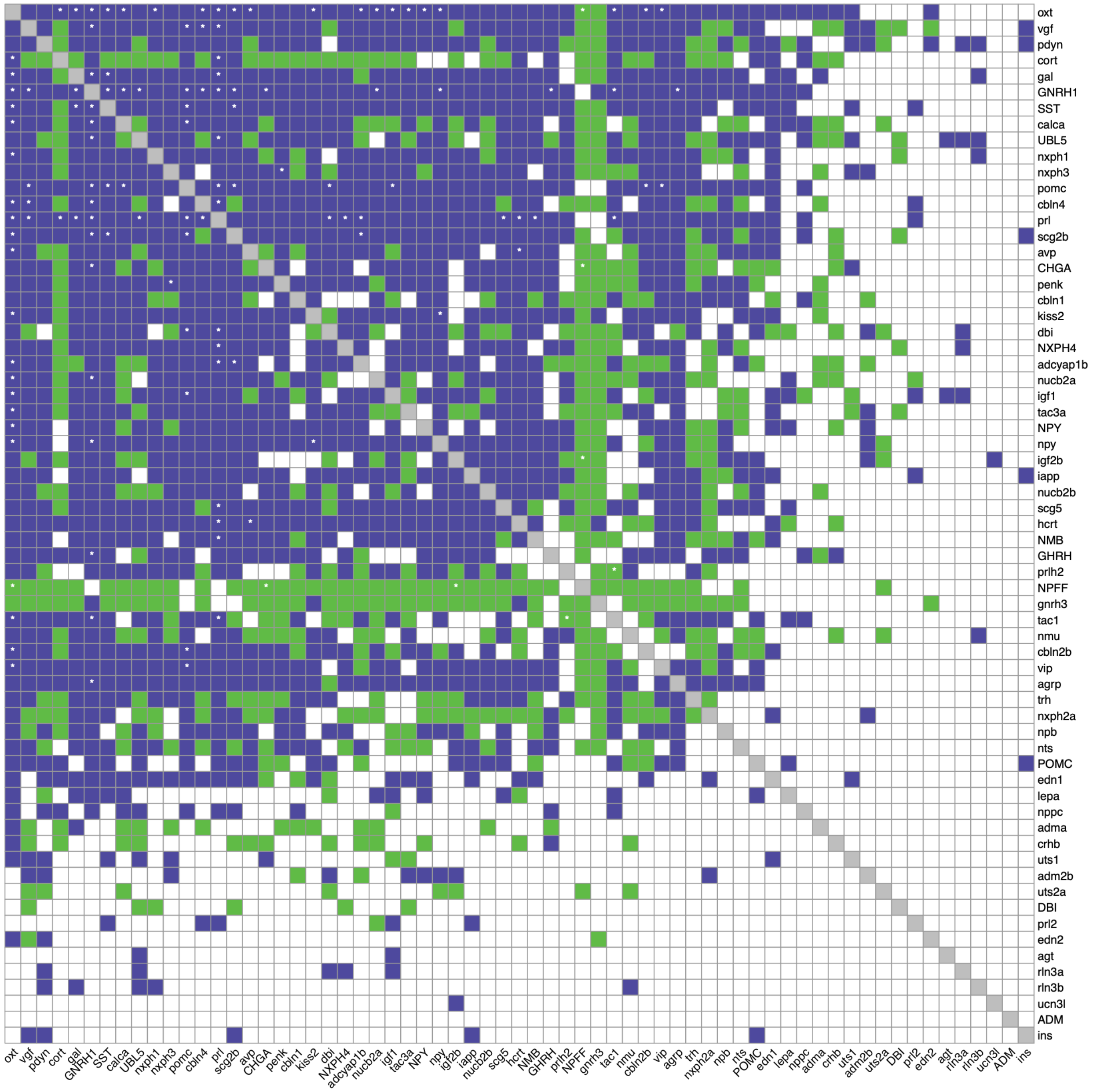


**Supplemental Figure 8.** Heatmap of difference between neuropeptide co-expression networks across social states (purple = increase in dominant, green = increase in subordinate, white = no difference) with significance (FDR pvalue < 0.15) desginated by white '*'.
